# Genome analysis and hyphal movement characterization of the hitchhiker endohyphal *Enterobacter* sp. from *Rhizoctonia solani*

**DOI:** 10.1101/2023.09.14.557762

**Authors:** Peiqi Zhang, Jose Huguet-Tapia, Zhao Peng, Ken Obasa, Anna K. Block, Frank F White

## Abstract

Bacterial-fungal interactions are pervasive in the rhizosphere. While an increasing number of endohyphal bacteria (EHB) have been identified, little is known about their ecology and impact on the associated fungal hosts and the surrounding environment. In this study, we characterized the genome of an *Enterobacter* sp. (En-Cren) isolated from the generalist fungal pathogen *Rhizoctonia solani*. Overall, the En-Cren genome size was typical for members of the genus and was capable of free-living growth. The genome was 4.6 MB in size, and no plasmids were detected. Several prophage regions and genomic islands were identified that harbor unique genes in comparison with phylogenetically closely related *Enterobacter* spp. Type VI secretion system and cyanate assimilation genes were identified from the bacterium, while common heavy metal resistance genes were absent. En-Cren contains the key genes for indole-3-acetic acid (IAA) and phenylacetic acid (PAA) biosynthesis, and produces IAA and PAA *in vitro*, which may impact the ecology or pathogenicity of the fungal pathogen *in vivo*. En-Cren was observed to move along hyphae of *R. solani* and on other basidiomycetes and ascomycetes in culture. The bacterial flagellum is essential for hyphal movement, while other pathways and genes may also be involved.

**Importance:** The genome characterization and comparative genomics analysis of En-Cren provided the foundation and resources for a better understanding of the ecology and evolution of this EHB in the rhizosphere. The ability to produce IAA and PAA may provide new angles to study the impact of phytohormones during the plant-pathogen interactions. The hitchhiking behavior of the bacterium on a diverse group of fungi, while inhibiting the growth of some others, revealed new areas of bacterial-fungal signaling and interaction yet to be explored.

## Introduction

Studies of the soil microbiome utilizing metagenomic tools have uncovered novel fungi and bacterial associations in the rhizosphere. The composition of these members is dynamic, influenced by the crop species and growth stage [1]–[4], soil type and nutrient levels [5]–[7] and other microbiome species [8], [9], indicating the possible existence of intricate relationships between microbes, crops, and the soil environment. The interaction between bacteria and fungi in the rhizosphere is likely to have profound impacts on plant biology and soil ecology [10]–[13]. Bacterial species can interact with fungal partners both externally and internally. Fungal-associated bacteria in the rhizosphere can affect fungal spore germination, and mycelium growth, provide a competitive advantage towards certain fungal species against another [14]–[16], improve soil conduciveness, and impact branching of the plant root system [17]. Endohyphal bacteria (EHB) can have profound roles in the ecology of the fungal hosts and the rhizosphere environment. Abundant *Burkholderia*-related Betaproteobacteria and Mycoplasma-related mollicutes are present in arbuscular mycorrhizal (AM) fungi in the phylum Glomeromycota (Bonfante & Desiro, 2017; Lastovetsky et al., 2018; Naumann et al., 2010; Spatafora et al., 2016). Alphaproteobacteria, gammaproteobacteria, and Firmicutes were identified within a wide range of ectomycorrhizal (ECM) and orchid mycorrhizal fungi in the phylum Basidiomycota (Bertaux et al., 2003; Deveau et al., 2010; Glaeser et al., 2016; Sharma et al., 2008). EHB has also been associated with fungal soil saprotrophs [18], [19] and plant pathogens [20]–[24]. *Mycetohabitans rhizoxinica* (formerly *Paraburkholderia rhizoxinica* or *Burkholderia rhizoxinica*) was identified from the rice pathogen *Rhizopus microspores* and controls the production of the plant toxins rhizoxin and rhizonin [22], [25].

The evolutionary implications and functional interactions between many identified EHB and their fungal hosts are yet to be established. Some obligate EHB that are associated with AM fungi are reliant on their fungal host metabolite pathways for essential carbon or mineral compounds and are marked by reduced genome sizes [18], [26]–[31]. Facultative symbiotic EHB, on the other hand, can exist as free-living bacteria when environmental conditions change; and genome reduction has not been observed [24], [32]–[34].

A strain of the *Enterobacter* genus, named *Enterobacter* sp. Crenshaw (hereafter En-Cren) was isolated from the hyphae of the soilborne, plant pathogenic fungus *Rhizoctonia solani* AG2-2IIIB, the causal agent of brown patch disease in cool-season turfgrass [20]. *Rhizoctonia solani* (teleomorph: *Thanatephorus cucumeris*) is a basidiomycete fungus with a broad host range causing sheath blight, black scurf, bare patch, root, and stem rot diseases on more than 200 plant species [35], [36]. On cool-season grasses, including bentgrasses, bluegrasses, fescues, and ryegrasses, *R. solani* AG2-2IIIB causes brown or tan patches of infected turf [37]. *R. solani* does not produce asexual spores and only occasionally form sexual spores. In nature, *R. solani* exists primarily as mycelia and survives as sclerotia in crop residues and soil [38]. The virulence factors of *Rhizoctonia solani* are largely unknown due to the lack of genetic manipulation studies with this species, in general. Previous studies have proposed various toxins, cell-wall-degrading enzymes (CWDEs), phenylacetic acid (PAA) and PAA derivatives, etc. [39]–[41]. The fungus showed less virulence when the EHB was cured from the fungus [20]. At the same time, the cured fungus produced less phenylacetic acid (PAA) than the wild-type fungus [20]. The EHB En-Cren was also observed to form a tight association with fungal hyphae, particularly movement to hyphal tips during the external growth of the bacteria [20].

In this study, we sequenced and annotated the En-Cren genome and conducted a genetic analysis of En-Cren regarding hyphal associated mobility and PAA production. Targeted and random mutations were conducted to generate En-Cren mutants to understand the requirements of PAA production and movement on fungal hyphal.

## Results

### The En-Cren Genome

En-Cren genome was sequenced with Oxford Nanopore Technology (ONT) MinION R9.4 flow cell and 2x250bp Illumina MiSeq. ONT sequencing resulted in 409,141 reads with a read length N50 of 19,058 bases and 4.13 GB total data, with approximately 890x genome coverage. The raw sequencing data length and quality were analyzed by Nanoplot (Version: 1.30.1) [42] and summarized (Table S1). The ONT reads were used to produce a draft assembly. Illumina MiSeq produced 984,179 pair-end reads, yielding 458.19 Mb total data and approximately 99x coverage. Illumina raw reads were evaluated with FastQC [43] and Nanoplot (Table S2) and used for polishing the draft assembly. A 4,632,138 bp single contig consensus genome was generated, following the Trycycler (Version: 0.4.2) workflow [44] and polishing with Medaka (Version: 1.2.1; © 2018-Oxford Nanopore Technologies Ltd.) and Pilon (Version: 1.24) [45]. The final assembly was circularized with Circlator (Version: 1.5.5) [46]. The assembly (NCBI GenBank database, GCA_024734705.1) has a total of 4272 coding genes as annotated by the NCBI RefSeq pipeline (Haft et al., 2018), 83 transfer RNA coding genes, and 25 ribosomal RNA coding genes (Table 1, Fig. 1).

**FIG 1.**
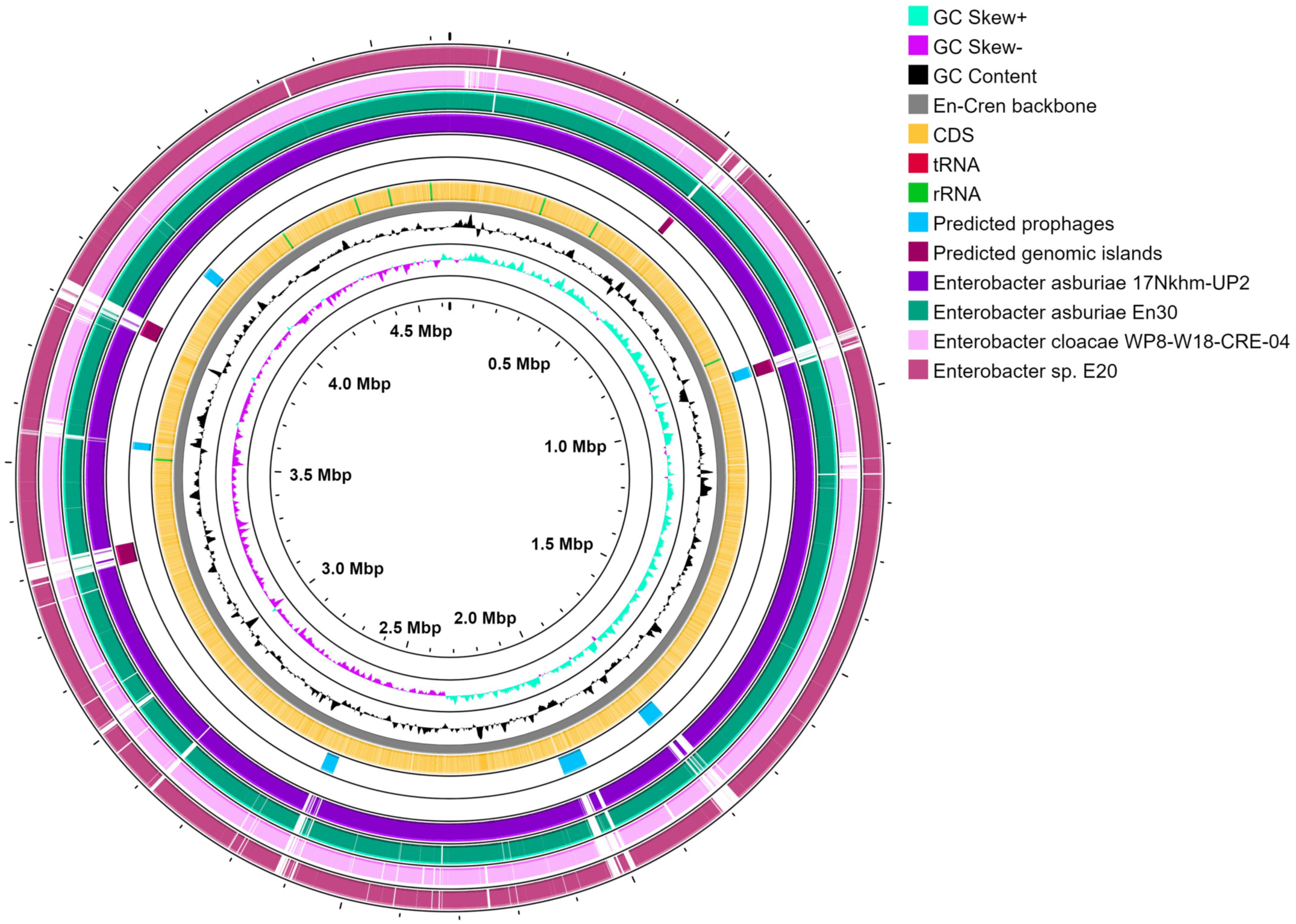
Circular view of En-Cren genomic feature annotations and comparison to four closest related *Enterobacter* isolate assemblies. From inside outwards, tracks showing genome position, GC skews, GC content, genome features including CDS and RNA, the six predicted prophage regions in En-Cren (with PHASTER), the predicted genomic island regions in En-Cren (with IslandViewer4), and sequence Blastn against related isolates *E. asburiae* 17khm-UP2, *E. asburiae* En30, *E. cloacae* WP8-W18-CRE-04, and *E.* sp. E20. Gaps indicate lack of corresponding sequence in related isolates.

**Table 1.** Genomic features of assembled En-Cren genome and closely related *Enterobacter* strains. En-Cren genome was annotated using the NCBI Prokaryotic Genome Annotation Pipeline (PGAP). Genomic features of other *Enterobacter* isolates were also obtained from the NCBI assembly database.

| Genome features | EnCren<br>(GCA_024734705.1) | *E. asburiae* 17Nkhm-UP2<br>(GCA_007035805.1) | *E. asburiae* En30<br>(GCA_017347425.1) | *E. cloacae* WP8-W18-CRE-04<br>(GCA_014169635.1) | *E. sp.* E20<br>(GCA_000801755.2) |
| --- | --- | --- | --- | --- | --- |
| Total genome size (bp) | 4,632,138 | 4,768,325 | 5,091,405 | 4,838,653 | 4,763,114 |
| Total GC content | 56.04 | 55.79 | 55.31 | 55.42 | 55.75 |
| Number of total genes | 4,414 (NCBI) | 4,559 (NCBI) | 4,981 (NCBI) | 4,787 (NCBI) | 4,551 (NCBI) |
| Numbers of coding genes | 4,272 (NCBI)<br>4,242 (Prokka) | 4,383 (NCBI)<br>4,367 (Prokka) | 4,796 (NCBI)<br>4,771 (Prokka) | 4,533 (NCBI)<br>4,628 (Prokka) | 4,371 (NCBI)<br>4,372 (Prokka) |
| Gene density | 0.915 genes per kb (1091 bases per gene) | 0.915 genes per kb (1091 bases per gene) | 0.937 genes per kb (1067 bases per gene) | 0.956 genes per kb (1045 bases per gene) | 0.917 genes per kb (1089 bases per gene) |
| Average CDS size | 972 | 968 | 942 | 924 | 966 |
| Number of transfer RNAs | 83 (NCBI)<br>84 (Prokka) | 83 (NCBI)<br>84 (Prokka) | 83 (NCBI)<br>86 (Prokka) | 86 (NCBI)<br>87 (Prokka) | 83 (NCBI)<br>85 (Prokka) |
| Number of ribosomal RNAs | 25 | 25 | 25 | 25 | 25 |
| Coding percentage | 89.0 | 88.7 | 88.3 | 88.3 | 88.6 |

### Phylogenetic Position of En-Cren in the *Enterobacter* Genus

A phylogenetic analysis of En-Cren was performed after screening the 321 available (as of June 2022) whole-genome assemblies of *Enterobacter* strains across all species in the NCBI genome database [47]. Thirty-two strains with more than 95% average nucleotide identity (ANI) to En-Cren, along with the type-strain of Enterobacter, *Enterobacter cloacae* subsp. *cloacae* ATCC 13047, were selected to construct a phylogenetic tree. The core genome analysis and alignment were conducted using Roary (Version: 3.12.0) (Page et al., 2015) (Table S3), and identified 2714 hard-core genes (100% presence), which was used to generate a maximum likelihood tree using RAxML (Version: 8.2.10) [48] with 100 bootstrap and GTR gamma model 9 (Fig. 2).

**FIG 2.**
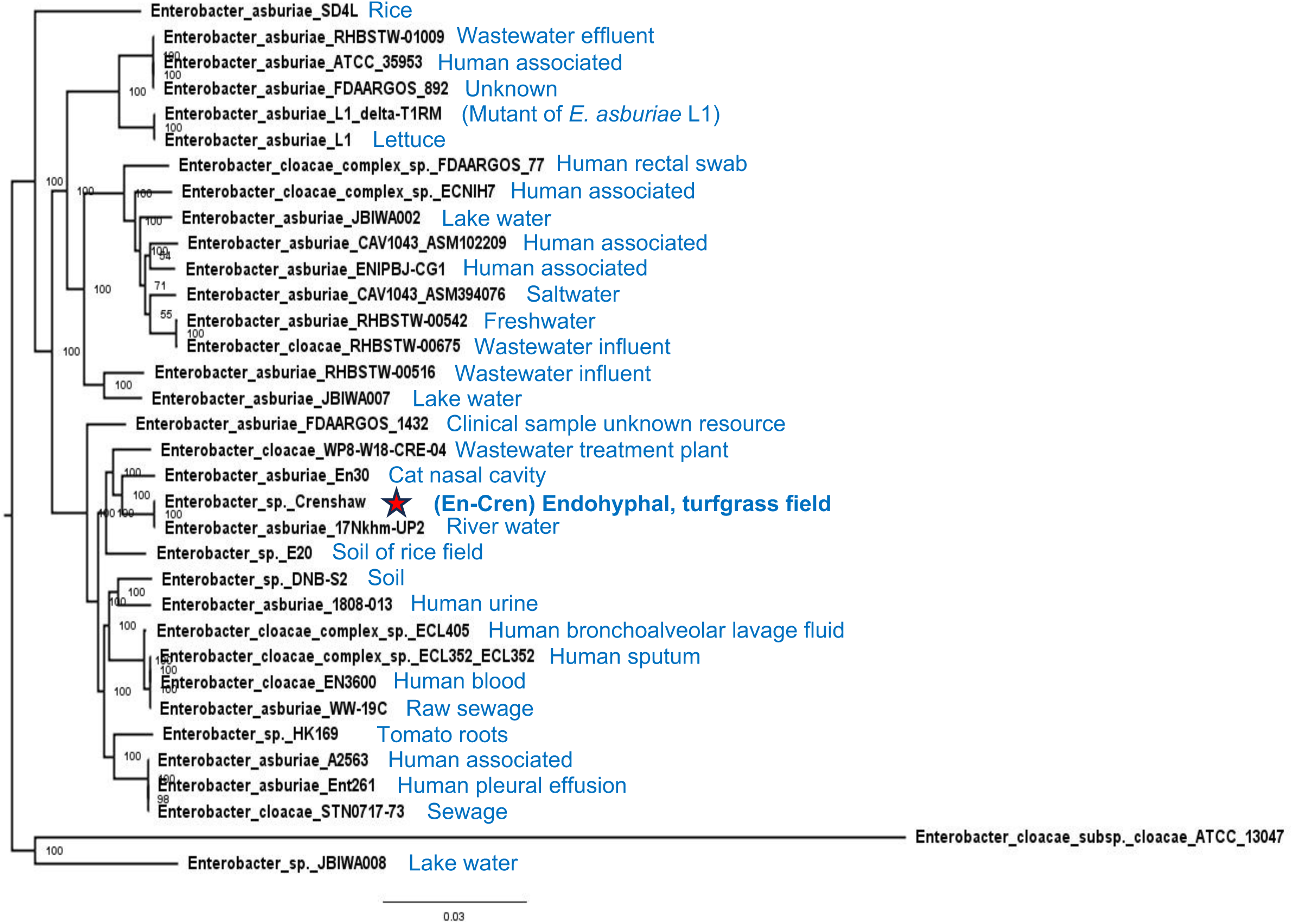
Core genome phylogenetic tree of En-Cren with 32 closely related isolates and *E.cloacae* type strain as an outgroup. 2714 core genes identified with Roary were used to generate a maximum likelihood tree using RaxML with 100 bootstrp and GTR gamma model. Node displaying bootstrap value. The source of the isolate for each genome assembly was labeled next to the name, genome accession numbers can be found in Table S7.

En-Cren is most closely related to isolates of *E. asburiae*, a few *E. cloacae* isolates, and a few unclassified environmental *Enterobacter* isolates, including *E. asburiae* 17Nkhm-UP2 (GenBank assembly: GCA_007035805.1), *E. asburiae* En30 (GCA_017347425.1), *E. cloacae* WP8-W18-CRE-04 (GCA_014169635.1) and *E*. sp. E20 (GCA_000801755.2). The origins of closely related isolates include river water, wastewater, glyphosate polluted soil of rice fields, and cat nasal cavity. The GenBank accession number and the corresponding isolate environment were summarized in Table S4.

### Comparative Genomics with Closely Related Strains

The genome size and features of En-Cren and the four most closely related isolates are summarized in Table 1. En-Cren has a smaller genome, higher GC content, and coding percentage than the other isolates. Gene-presence-absence analysis on En-Cren and the thirty-two isolates with higher than 95% ANI identified unique genes, many encoding uncharacterized hypothetical proteins (Table S5). Notable among the annotated unique genes, En-Cren genome contains genes related to a *cynTSX* cyanase operon [49](NWV12_18030, NWV12_18035, with the regulator gene *cynR* NWV12_18025, and the *cynX* encoding the transporter NWV12_08375), which enables degradation and utilization of cyanate as a nitrogen source. These genes were located on a genomic island and could be the result of horizontal gene transfer. Homologs for En-Cren *cynS*, a gene encoding the cyanate hydratase, or cyanase, were identified only in a small number of strains within the *Enterobacter* genus, while also present in some strains of *Pseudomonas*, *Dickeya,* and *Kosakonia*. Meanwhile, En-Cren lacks some genes present in more than half of the closely related thirty-three isolates, noticeably ompN_2 encoding outer membrane porin N (functionally resembles OmpC) [50], translocation and assembly module TAM subunit B (which is essential for virulence of several pathogenic bacteria) [51], copper-resistance genes and, sliver binding and exporting genes which are characteristic of some bacteria species inhabiting polluted agricultural soil and clinical environments [52]–[55] (Table S6).

### Prophage and Genomic Island Regions

The prophage identification tool PHASTER (PHAge Search Tool-Enhanced Release) was used to identify six prophage regions in the En-Cren genome (Table 2). Prophage region 6 was present in all closely related *Enterobacter* isolates (Fig. 1), while prophages 1, 3, and 5 all have >99% identical BLASTN hits with over 90% coverage with other *Enterobacter* spp. in the NCBI database. BLASTN search of prophage region 4 yielded fragmented alignments with a few species of *Enterobacter* for around 70-81% coverage and 93.6%-95.7% identity. Notably, prophage region 2 only had scrambled alignments with *Enterobacter* spp. for less than a total of 67% coverage. These prophage regions contain mainly genes encoding hypothetical proteins, phage-structure proteins, outer membrane lipoproteins (YnfC family lipoprotein, cor protein) (Arguijo-Hernández et al., 2018), lexA repressors and various DNA repair and homologous recombination proteins (recombinase RecT, PD-(D/E)XK nuclease-like domain-containing protein, replication proteins, DinI family protein, crossover junction endodeoxyribonuclease RusA, translesion error-prone DNA polymerase V autoproteolytic subunit, Y-family DNA polymerase, YnfU family zinc-binding protein). They also carry genes encoding stress-resistance related proteins including toxin/antitoxin PasT/PasI (preserved in many *Proteobacteria*) [56], antibiotic-resistant chloramphenicol acetyltransferase, cold shock adaptation proteins (kdo(2)-lipid IV(A) palmitoleoyltransferase LpxP, cold shock domain-containing protein, cold shock-like protein CspF). Proteins potentially act in host interactions were also present, including lysozymes, endolysins, enterohemolysin, and inhibitors of vertebrate lysozymes (Ivy), host immune evasion related O-acetyltransferase OatA [57], and host interaction and counter defensive related cyclic beta 1-2 glucan synthetase (Rigano et al., 2007). Most of the unique genes identified through comparative genomics analysis were found in these prophage regions (Table S5). During the culture of En-Cren mutants, plaque-like spots, likely caused by phage excision, were observed. Subsequent sequencing of a flagellar mutant strain *fliR-* revealed that the prophage region 4 was missing, indicating phage excision (Fig. S1).

**Table 2.**
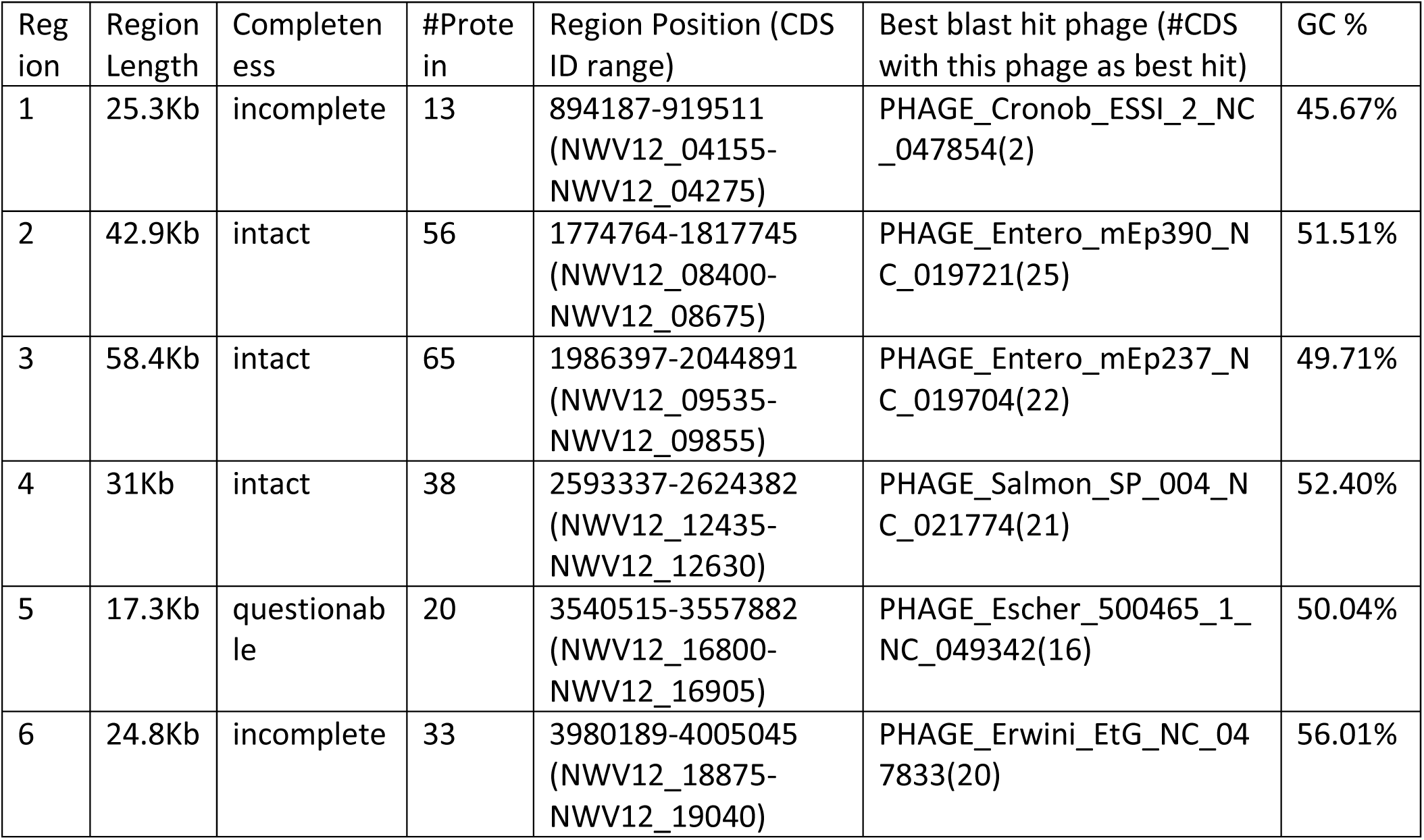
Prophages in EnCren genome as identified using PHASTER. Completeness of prophage regions characterized by PHASTER according to the presence of phage-related proteins and region sizes. Best blast hit phage identified by blasting each CDS within the region against the Virus and Prophage Database.

The En-Cren genome also contains several genomic islands other than the prophage regions (Fig. 1). These islands contain many hypothetical proteins, often with integrase genes next to tRNAs. These regions feature cyanate assimilation enzymes [49], vitamin biosynthesis and utilization enzymes (pyridoxamine 5’-phosphate oxidase family protein and acetolactate synthase large unit/ thiamine pyrophosphate enzyme) [58], [59], transcription regulators, transporter proteins, alkaline-shock proteins, chemotaxis proteins, and toxins (Table S7). Notably, in the genomic island region 3, En-Cren obtained an entire cluster of type VI secretion system (T6SS) proteins, which was absent in the four closely related *Enterobacter* strains, although present in some other *Enterobacter* spp. Most genes located on genomic island 4 and a few on islands 2 and 3 were absent from all other 32 related strains (Table S5).

### Chitinases and Chitin-related Genes in En-Cren

Two copies of the chitinase gene (NWV12_06925, NWV12_06935, EC 3.2.1.14, conserved among Enterobacterales) were located near the Type II secretion system operon in the En-Cren genome. The *chbBCARFG* operon (NWV12_08840-NWV12_08865) for uptake and metabolism of chitobiose (conserved in *Enterobacter* spp.) and a gene encoding chitoporin (NWV12_05940), which is important for chito-oligosaccharides utilization, were also present in the En-Cren genome. When cultured on 1% colloidal chitin YEM medium, En-Cren did not show clearing zones that indicate chitinolytic activity.

### PAA and IAA Production of En-Cren

En-Cren contains the gene *ipdC*, encoding indole-3-pyruvate decarboxylase, which has been previously shown to be involved in the biosynthesis of the plant growth regulators indole acetic acid (IAA) [60]–[62] and PAA [63]. Free-living En-Cren can produce PAA in ¼ strength potato dextrose broth (¼ PDB) medium supplemented with phenylalanine (Phe). Upon introduction of a mutation in *ipdC*, PAA production drops to low levels (Fig. 3A). When grown in the minimal liquid medium (MM, with citric acid as carbon source), En-Cren produced trace levels of PAA in comparison to growth in ¼ PDB, even with additional phenylalanine precursor (Fig. 3A). Complementation of the *ipdC* mutant of En-Cren restored the PAA production to the wild-type (WT) level in 1/4 PDB medium and not in MM.

**FIG 3.**
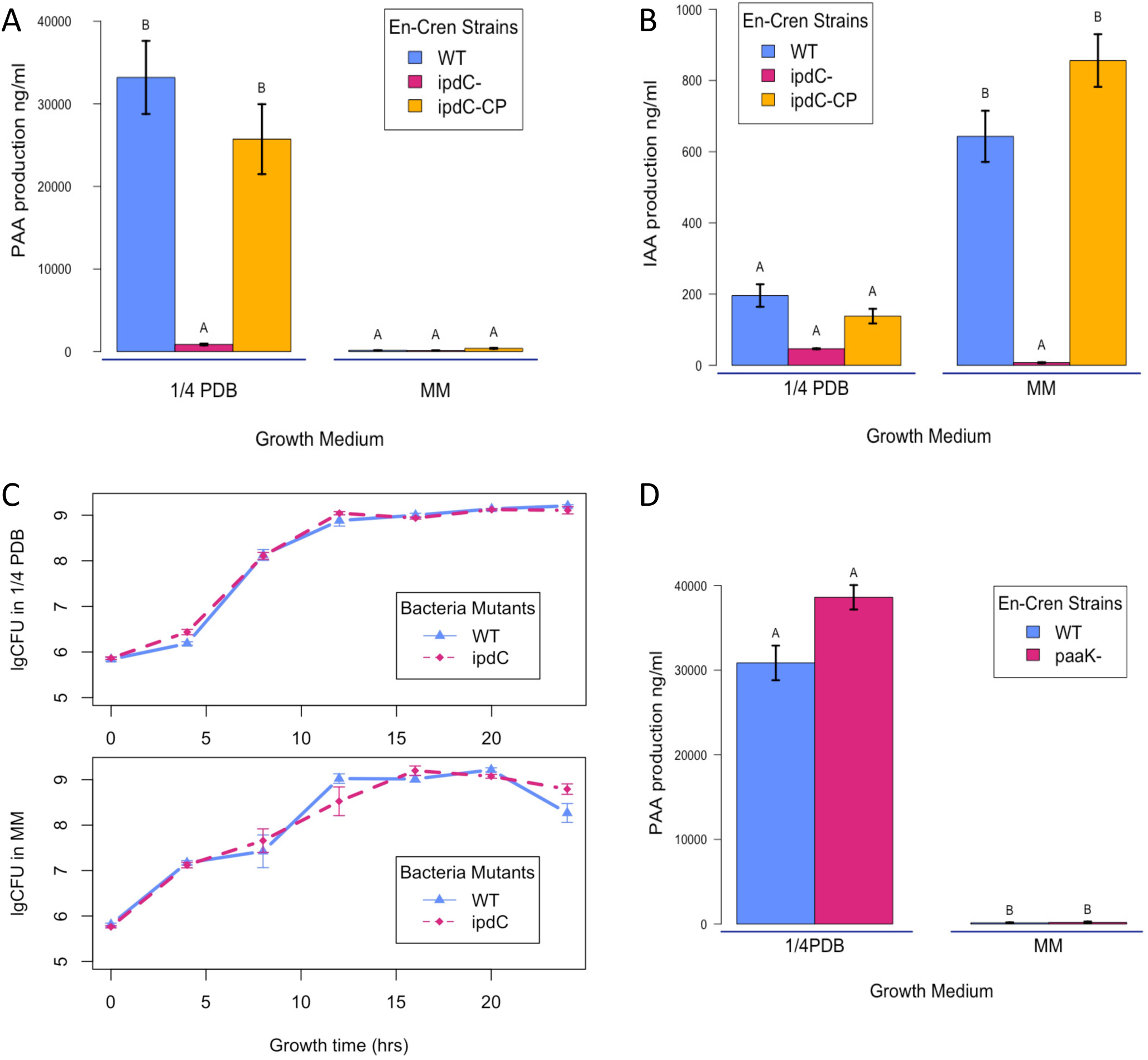
PAA and IAA production of En-Cren bacteria wildtype (WT) and mutants. (A) PAA production of free-living En-Cren bacterium WT, *ipdC-* mutant and *ipdC-* mutant complementation strain (*ipdC-CP*) in ¼ PDB and MM (+ citric acid) liquid medium supplemented with 1% phenylalanine. (B) IAA production of free-living En-Cren bacterium WT, *ipdC-* and *ipdC-CP* in ¼ PDB and MM (+ citric acid) liquid medium supplemented with 1% L-tryptophan. (C) Bacterial growth curve of WT and *ipdc-* En-Cren in quarter strength PDB and minimal media (+citric acid). (D) PAA production of En-Cren WT, *paaK-* mutant in ¼ PDB and MM (+ citric acid) liquid medium supplemented with 1% phenylalanine.

En-Cren can produce IAA in both growth media with precursor L-tryptophan added, and the *ipdC* mutant reduced the levels of IAA. Complemented strains were able to produce IAA at the same level as the WT strain (Fig. 3B). The growth rate of WT and *ipdC*-mutant En-Cren is similar in both growth media (Fig. 3C).

The lack of PAA presence in MM has prompted us to explore whether PAA was degraded in MM condition. Mutant of the *paaK* gene, encoding enzyme phenylacetyl-CoA ligase for the first step of PAA catabolism [64], [65], was created and tested for PAA production in both media. The *paaK-* mutant did not show an increase in PAA production when grown in ¼ PDB or MM compared to WT En-Cren (Fig. 3D).

### En-Cren Movement on Fungal Hyphae

Upon contact with hyphae, En-Cren moves along the host *Rhizoctonia solani* isolate (named Rs-Cren) hyphae and associates closely with the fungal culture. The plasmid pVIO1-2, which contains genes for violacein production and gives the bacteria purple pigmentation, was added to En-Cren to enhance visualization. Rs-Cren hyphae-agar plug was placed at the center of ¼ strength LB culture plate, and the WT En-Cren with violacein producing plasmid pVIO1-2-Tet (WT-pVio) were streaked on the same agar plate 1 cm from the hyphal plug. Upon contact, bacteria migrated along the hyphae of Rs-Cren, covering the mycelia in about two days (Fig. 4A). A video made from 96-hour time-lapsed photos also shows the progress of En-Cren movement on Rs-Cren mycelia. (MP4 file, 28M. https://ufdc.ufl.edu/IR00011445/00001.)

**FIG 4.**
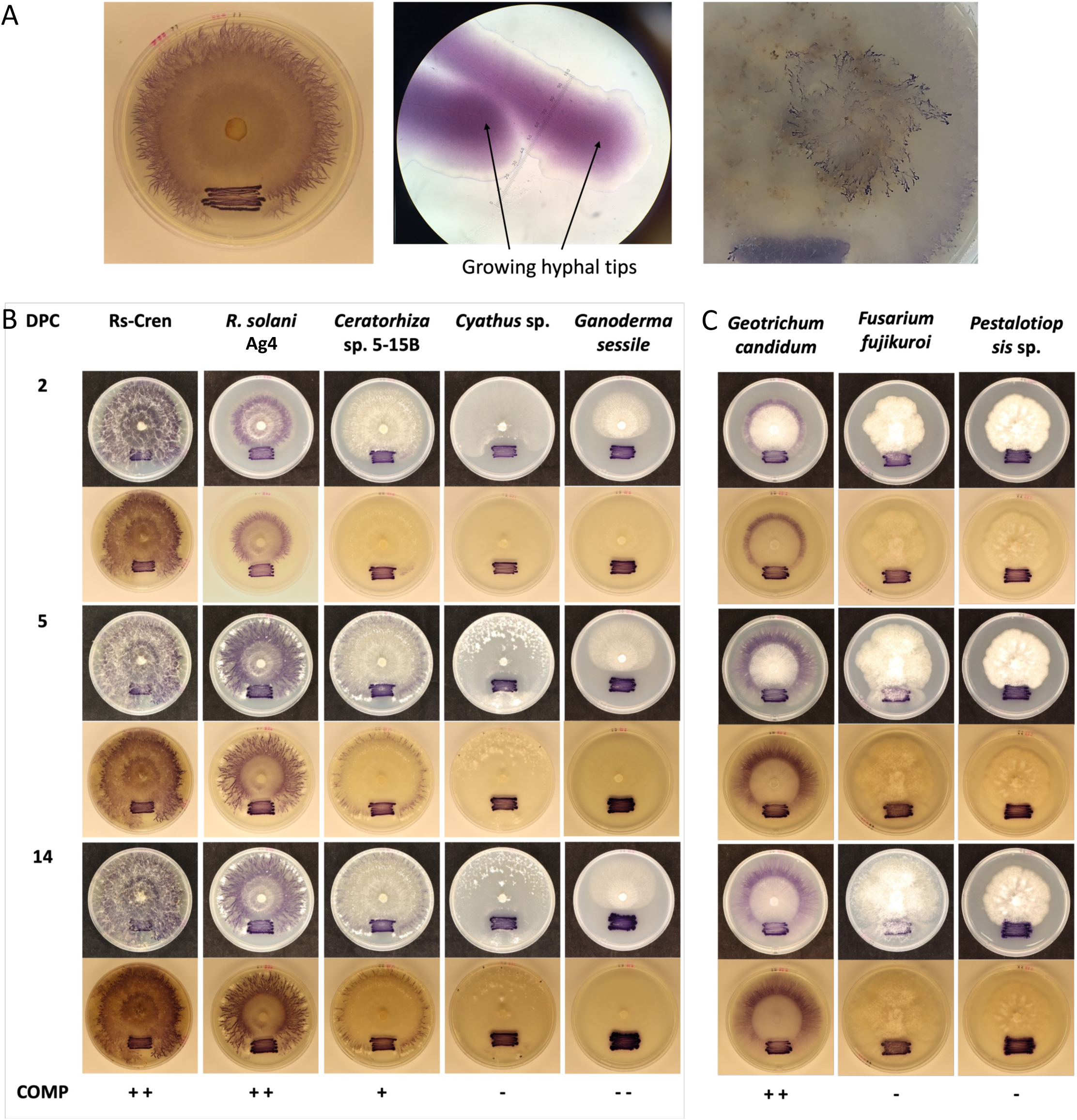
En-Cren movement and aggregation along fungal hyphal when co-cultured with different fungal cultures on ¼ LBA plates. (A) En-Cren movement and aggregation along Rs-Cren hyphae, including on the aerial hyphae. Movement along aerial hyphae was tested by preparing medium on the lid of petri-dish and incubate the co-culture plates face-up. (B) En-Cren movement along hyphae of other *Basidiomycete* isolates including another *R. solani*, *Ceratorhiza* sp., *Cyathus* sp. and *Ganoderma sessile*, compared to movement on host fungus Rs-Cren. Photo taken 2, 5 and 14 Days after the growing periphery of the fungus approached the bacterial inoculation site. (C) En-Cren movement along hyphae of some *Ascomycete* isolates including *Geotrichum candidum*, *Fusarium fujikuroi* and *Pestalotiopsis* sp., compared to movement on host fungus Rs-Cren. Photo taken 2, 5 and 14 Days after the growing periphery of the fungus approached the bacterial inoculation site.

En-Cren movement along Rs-Cren aerial hyphae was also observed, as shown by En-Cren WT-pVio bacteria growth on the lid of the co-culture plates (Fig. 4A). The hyphal movement of En-Cren on different basidiomycete (Fig. 4B) and ascomycete fungi (Fig. 4C) was also tested. A similar hyphal movement pattern was observed on different strains of *Rhizoctonia solani*, with En-Cren WT-pVio spreading in two days after contact with a lab strain of *Rhizoctonia solani* Ag4. After 5 days of contact, visible bacteria migrated toward the hyphal tips of the *Rhizoctonia*-related fungus, *Ceratorhiza* sp. No bacterial growth was observed around the center of the fungal colony. En-Cren WT-pVio showed some inhibitory effect on *Cyathus* sp. and *Ganoderma sessile*. The inhibitory effect was also observed with En-Cren WT without the pVio plasmid (Fig. S2). After 14 days of co-culture, En-Cren WT-pVio was observed migrating along the hyphae of *Cyathus* sp. towards the bottom of the plate. No movement was observed with *G. sessile*.

En-Cren WT-pVio showed migration on the ascomycete *Geotrichum candidum* two days after contact (Fig. 4C). No visible bacterial movement was observed after 14 days of co-incubation on *Fusarium fujikuroi* and a *Pestalotiopsis* sp. Notably, growth of the *Pestalotiopsis* sp. strain was slow on ¼-strength LB medium. When tested on a *Sclerotinia sclerotium* strain, visible En-Cren hyphae migration was only observed starting 3 days post contact and still limited to adjacent hyphae after 5 days (Fig. S3).

### Genes Affecting Bacterial Movement along Hyphae

Motility mutants of En-Cren were constructed, targeting flagellar motility and the type IV pili motility to determine the factors involved in the hyphal movement. Targeted mutations were created in En-Cren genes *flgE* encoding the flagellar hook protein, *flgD* encoding the flagellar basal-body rod modification protein, *flgF* encoding flagellar basal-body rod protein, *motA* encoding flagellar motor stator element, *fliC* encoding flagellin protein, *flhA* encoding the flagellar biosynthesis protein involved in the export of flagellum proteins. These flagellar mutants all have impaired hyphal movement on Rs-Cren culture. Meanwhile, the swimming motility ability of these flagellar mutants was demonstrated to be impaired on 0.3% agar ¼ LB plates. No swarming phenotypes were observed with En-Cren strains under tested agar concentrations (0.5% and 1.0%). The growth rates of these flagellar motility mutants did not differ significantly from the WT En-Cren (Fig. S4).

*pilT* mutants of the type IV pili (encoding the type IV pilus retraction ATPase) retained the ability to spread along the fungal hyphae, but the bacteria were only observed to cover the whole fungal colony after 5-7 days compared to 2 days for WT En-Cren. Other type IV pilus mutants, including *pilQ, pilC, and pilA* (encoding the type IV pilus secretin, assembly protein, and major pilin protein), showed a similar pattern to the WT En-Cren strain. The swimming motility of the pili mutants, except for *pilT-*, was not affected (Fig. 5).

**FIG 5.**
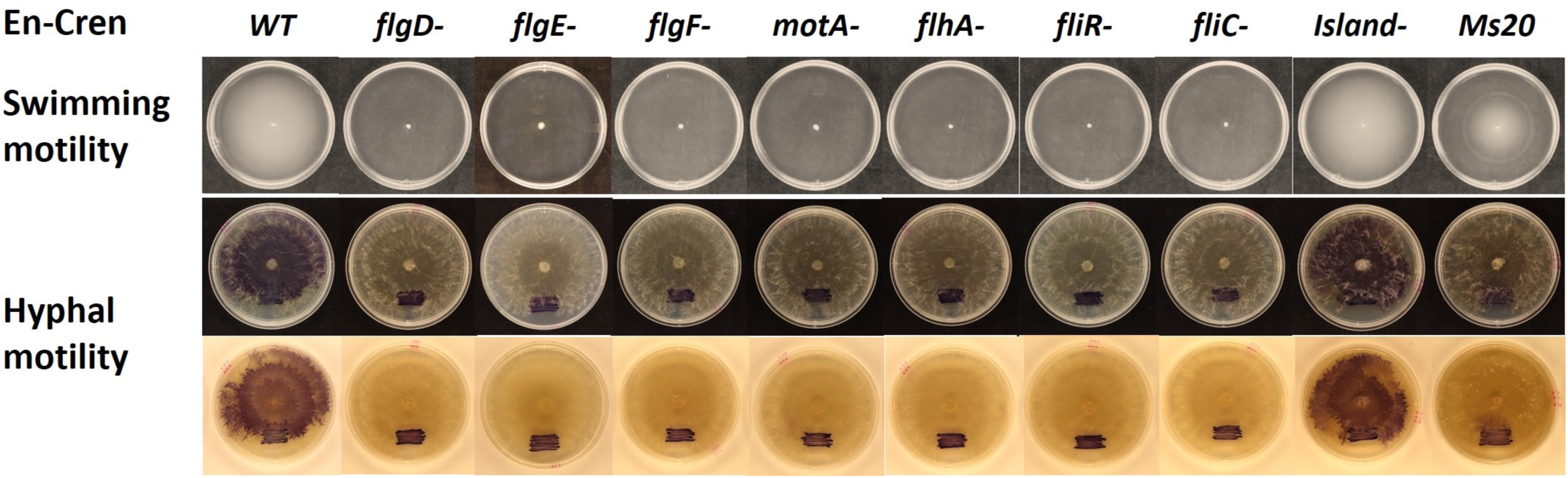
The swimming motility and fungal hyphae movement feature of En-Cren WT and mutants. Swimming motility of bacteria was tested on ¼ strength LB soft agar plates, after 12 hours of inocubation at 30 °C. Hyphal motility patterns of En-Cren WT and mutants on co-cultured Rs-Cren were captured and compared 2 days after the growing periphery of the fungus approached the bacterial inoculation site.

In addition to direct mutagenesis to flagella and type IV pili related genes, we conducted random mutagenesis of the En-Cren with transposon Tn5 kit EZ-Tn<KN>. Bacterial mutant libraries with transposon insertion at random genome locations were generated and screened for hyphal movement. Individual mutant colonies that show no hyphal migration upon contact with a growing fungal colony were selected for further testing. We have identified hyphal-movement impaired mutants that have insertion at genes *fliR*, *flhC*, *flhD,* and *phdR* (Table S8). Among the identified mutants, *fliR*, *flhC*, *flhD* are genes encoding protein components of the flagella and were also impaired for swimming motility. The *pdhR* gene encodes the pyruvate dehydrogenase complex repressor and is the first gene in the operon encoding proteins comprising the pyruvate dehydrogenase complex. Pyruvate dehydrogenase is known to control the glycolysis metabolic and citric acid cycles.

## Discussion

Characterization of En-Cren resulted in a genome size of 4,632,138 bp. In contrast to other *Enterobacter* genomes, En-Cren does not display an apparent genome reduction sometimes associated with obligate endohyphal bacteria species. [33]. Combined with the observation that En-Cren can also be free-living outside the fungal host, the results indicate that En-Cren is a non-obligate symbiont. Although most identified closely related strains were named *E. asburiae*, we did not adopt the species name due to the confusion in the *Enterobacter* complex taxonomy. The phylogenetic placement of En-Cren with some environmental strains indicates that En-Cren could occupy a wide range of niches and that related *Enterobacter* strains isolated from environmental samples may also have the potential to form associations with fungal species in the rhizosphere. *Enterobacter* spp. endophytes, phytopathogen, and animal gut commensals are associated with various natural habitats. In contrast, only certain strains of some species/subspecies were identified as opportunistic nosocomial pathogens due to their antibiotic resistance features [66], [67]. Accordingly, we found that identified strains with more than 95% ANI to En-Cren have various sources of origin, including environmental, clinical, and some plant samples. As most studies are focused on clinical isolates rather than plant associated and soil and water samples, En-Cren and many related *Enterobacter* species may be more prevalent in various natural environments. We have identified unique prophage regions and other genomic islands from En-Cren encoding a Type VI secretion system, cyanate assimilation genes, stress-response proteins, host immunity related proteins, transcription regulators, transporter proteins, and toxins. Meanwhile, some heavy metal resistance genes commonly found in soil inhabiting *Enterobacter* spp. were absent. These differences in the stress regulation and metabolite arsenal of En-Cren could be evidence that En-Cren faces different environmental stresses than other *Enterobacter* isolates. A closer look into these proteins may grant us more hints about En-Cren’s niche adaptation.

En-Cren has genes encoding for chitinase and T2SS-related genes, which were also identified to be critical for fungal infection process in the *M. rhizoxinica* interaction [28], [68]. En-Cren also contains a gene encoding chitoporin with secretion signal peptide, which is identified as responsible for chito-oligosaccharides transportation and uptake in the marine bacterium *Vibrio harveyi* [69], [70]. Bacterial chitinases are often associated with a potential role in fungal/insect antagonism and biocontrol [71]. Some strains of the *Enterobacter* genus, including *E. cloacea* and *E. agglomerans* isolated from soil samples, have been described to exhibit strong chitinolytic activities [72]–[74] and have shown antifungal activities towards fungal plant pathogens, including *R. solani* [72]. However, differences in the chitinolytic abilities were often observed for different strains of the same bacterial species, even in strains identified from the same environment [75], [76]. Although free-living En-Cren did not show chitinolytic behavior when cultured on colloidal chitin YEM medium, further testing of the chitinolytic ability of En-Cren under different nutrient and pH and under fungal co-culture conditions may reveal conditions where En-Cren utilizes chitinases. Similar to most sequenced *Enterobacter* spp., En-Cren does not contain a T3SS, which is an important bacterial secretion system for many pathogenic bacteria [77]–[79]. T3SS has been shown essential for the EHB *M. rhizoxinica* to be reintroduced into the fungal host and induce host sporulation [80], and secreting TAL effector-like proteins that influence fungal host transcription and stress tolerance [81]. Notably, different strains of EHB *M. cysteinexigens*, which were isolated from non-pathogenic fungus *M. elongata*, possess different putative T3SSs [82]. The lack of T3SS in the En-Cren genome indicates that En-Cren utilizes different invasion and signaling pathways for host interaction. Notably, En-Cren contains genes for a set of T6SS, which is absent from the most closely related Enterobacter strains. Some other EHB, namely the above-mentioned *M. cysteinexigens* B1-EB^T^, and a *Burkholderia* sp. 9120 strain identified from foliar fungal endophytes, also possess the T6SS [32], [82]. Although T6SS has been mainly shown to function in bacterial-bacterial interaction and, in some instances, host-microbial interaction [83], the importance of the system in endohyphal bacteria still awaits investigation. In a recent dual RNA-seq analysis of EHB *Luteibacter* sp. cocultured with its host foliar endophyte fungus *Pestalotiopsis* sp., 11 of 13 of the predicted bacterial T6SS genes were upregulated compared to the axenic free-living bacterial culture [84]. This further supports the hypothesis that T6SS may play a role in the establishment of bacterial-fungal symbiosis relationship.

Some other genes previously shown to be involved in EHB host interaction include vitamin B1 and B12 biosynthesis genes [27], [58], [85], gene encoding O-antigen of the bacterial outer membrane [86] and lipopeptides [87]. En-Cren also contains homologous genes for B1 and B12 biosynthesis and transportation pathways and a gene cluster for lipopolysaccharide biosynthesis proteins, an O-antigen ligase, and glycosyltransferases. Recent studies on the function of vitamin B6 in the oxidative stress regulation and disease development of *Rhizoctonia solani* are also worth noting [88], [89], especially as we also identified an En-Cren unique gene related to vitamin B6 oxidation. A transcriptomic study on free-living and fungus-associated En-Cren on different nutrient media may identify differentially expressed bacterial genes for fungal surface colonization or invasion and provide more comprehensive insight into pathways involved in bacterial-fungal interactions.

Bacterial biosynthesis of auxin and the proposed roles played by microbial auxin production was also an area of interest in this study, as previous studies present potential plant growth and disease control benefits from microbially produced IAA and PAA [63], [90]–[93]. The possibility that biosynthesis of PAA by *R. solani* may be associated with endohyphal bacteria motivated an analysis of PAA biosynthesis by En-Cren. Although the pathways for PAA biosynthesis in bacteria have not been fully elucidated, the En-Cren *ipdC* gene (encoding a pyruvate decarboxylase/ alpha-keto decarboxylases) functions in both IAA and PAA biosynthesis, as in the case of *E. coli* and *A. brasilense* [63], [94], [95]. Loss of function resulted in low PAA or IAA levels by free-living bacteria, depending on culture conditions and supplementation with the respective precursors. Since En-Cren IAA production in both growth media demonstrated that the *ipdC* gene is still functional under MM condition, the lack of PAA presence in MM is likely due to other reasons, including restricted uptake of phenylalanine, additional regulators for PAA synthesis, or degradation of PAA. As phenylacetate-CoA ligase encoded by *paa*K gene is the enzyme catalyzing the first step of the bacterial PAA catabolite pathway [96], and our *paaK*-mutant En-Cren produced a similar trace level of PAA in the MM, it is unlikely that PAA degradation was the cause. Mutants of genes encoding other enzymes in the PAA biosynthesis pathway, or in the resembled IAA biosynthesis pathway may help us further interpret this observation. Noticeably, the IAA production for WT strain En-Cren in ¼ PDB is significantly less than that in minimal medium. This difference in IAA production level may be due to nutrient and substrate variability, as documented before in *A. brasilence* [97]. Notably, the EHB *Luteibacter* sp. enhances the IAA production of the foliar endophyte *Pestalotiopsis* sp. aff. *neglecta* [90], and a recently isolated EHB *Klebsiella aerogenes* (Syn. Enterobacter aerogenes) of mycorrhizal fungus *Fusarium oxysporum* KB-3 was found to produce IAA and contribute to the plant growth promoting effect of its fungal host [98]. These findings raised the possibility that En-Cren IAA biosynthesis may play a role in the interactions with *R. solani* and its plant host. Previous experiments indicated enhanced PAA production in En-Cren inhabiting *R. solani* Rs-Cren [20]. PAA has also been shown to have antifungal activity on *R. solani* when produced by *Streptomyces humidus* bacteria [99] and suppression of spore germination of *Fusarium oxysporum* when produced by a strain of *Bacillus mycoides* [100]. Understanding the effects of IAA and PAA biosynthesis by WT En-Cren on the physiology and virulence of the fungal pathogen Rs-Cren will be useful in defining the role played by IAA and PAA in *R. solani* virulence.

En-Cren can migrate along the hyphae of not only its host fungus *Rhizoctonia solani* strain Rs-Cren, but also another isolate of *R. solani* in a different anastomosis group, an isolate of a closely related basidiomycete *Ceratorhiza* sp., and an isolate of a phylogenetically distant ascomycete *Geotrichum candidum* sp. This ability to migrate on mycelia of diverse fungi has not been documented in *Enterobacter* sp. previously but has been noted for *Burkholderia terrae* [101] and *Serratia marcescens* [19]. In contrast, En-Cren showed no movement on *Ganoderma sessile*, *Fusarium fujikuroi,* and *Pestalotiopsis* sp., while even causing growth inhibition towards *G. sessile*. Species-specific movement patterns were also observed with hyphal migrating bacteria *B. terrae* [101] and *S. marcescens* [19] when tested on fungi from different phyla. This difference in the movement patterns is possibly due to specific chemotaxis events, including nutrient attraction and toxin compelling. The ability of En-Cren to move along the fast-growing tips of the hyphae, combined with little observed migration and accumulation around aged Rs-Cren culture, may indicate that the compatible fungi are providing nutrients for the bacterium. In the case of *Pseudomonas putida,* the bacterium was attracted to the oomycete partner, similar to the known chemoattractant sodium salicylate [102]. Toxin or substrate repelling is also plausible, as supported by the growth inhibition effect towards *Cyathus* sp. and *Ganoderma sessile* by En-Cren during the co-culture. Alternative explanations include the antimicrobial potential of violacein, although evidence on its antifungal toxicity remains scarce and inconsistent [103], [104], and WT En-Cren without the violacein producing plasmid also showed a growth inhibition effect towards both fungi. Other supporting examples for the repelling compound theory include bacterium *S. marcescens* with ascomycete *Aspergillus fumigatus* [19]. As *S. marcescens* could not migrate on the living culture of *A. fumigatus* but was able to migrate and cover dead hyphae of this fungus, indicating that a toxin or repellent was secreted by living *A. fumigatus*.

En-Cren’s capability to migrate along fungal hyphae, including aerial hyphae (at least that of Rs- Cren), could provide survival and competitional advantages to the bacterium [105], [106]. When tested in simulated soil environments [19], [102], [107], bacteria could hitchhike and rapidly disperse on fungal mycelium through multi-layers of different material. This may present a way of bacterial dispersal in the natural environment, even across air pockets of unsaturated soils. As *Rhizoctonia solani* can utilize dead matter in soil [108], [109] and spread mycelium in porous soil structures to approach potential susceptible host [110], [111], it’s plausible that En-Cren would take advantage of this fungal highway. Mutations in En-Cren flagellar genes impaired the ability to move along fungal hyphae, supporting the role played by flagellar motility in fungal-bacterial interactions of En-Cren, as seen in some other bacterial species [19], [105], [107], [112]. The type IV twitching motility genes did not affect movement along hyphae, but the *pilT* gene mutant seemingly reduced the movement speed while also showing slower swimming motility in soft agar. It has been observed that flagellar mutants (including those in *fliA* and *flhD*) in *S. marcescens* only slowed down migration along hyphae [19] and that some flagellar positive bacteria closely related with mycelia migrators did not show hyphal movement ability [105]. Also, type I fimbriae mutants of *S. marcescens* tested have shown faster movement along fungal hyphae [19]. Since mutant screening only identified a few genes in the flagellar apparatus, it is likely screening has not saturated the available motility genes. Thus, we hypothesize that alternative modes of motility are involved and that further screening and targeted mutagenesis in genes, including adhesion related genes, could help us better understand the hyphal movement phenomenon.

Studying En-Cren/Rs-Cren interaction and bacterial-fungal interaction, in general, could lead to a better understanding of the ecology of many soil bacteria that could form closer than anticipated associations with various fungi and plants.

Many questions remain in the En-Cren/Rs-Cren interaction. When and how does En-Cren invade its fungal host? Can the bacterium produce an effective amount of PAA or IAA during natural setting to affect the virulence of the fungal host or the biology of the associated plant host? Can En-Cren survive within the fungal sclerotia for a prolonged period? What are the attractants or toxic compounds secreted during En-Cren interaction with different species of fungi? Further field testing of En-Cren’s survival rate and virulence contribution and transcriptomic analysis on En-Cren cultured with and without various fungal partners may help us find more clues.

## Methods and Materials

### Strains and Culturing of Bacteria and Fungi

The free-living En-Cren was isolated previously from *R. solani* strain Rs-cren [20]. Unless otherwise stated, the bacterium was cultured at 28°C on Luria-Bertani agar (LBA) plates or in Luria-Bertani broth (LB). Fungal strains with lighted-colored filamentous mycelium that belong to distinct genera of Ascomycota and Basidiomycota were obtained from the Fungal Teaching Lab at the Department of Plant Pathology, University of Florida, and are listed in Table S9. Fungal cultures were maintained on potato dextrose agar (PDA) medium, or PDA acidified (pH ∼4.8) with 1 ml of 85% lactic acid per liter (APDA) and incubated at room temperature unless otherwise stated.

### Bacterial DNA Extraction and Sequencing

Genomic DNA was extracted using the Wizard Genomic DNA Extraction Kit (Promega). DNA concentration and quality were determined by NanoDrop One Microvolume UV-Vis Spectrophotometer (Thermo Scientific). Sequencing of the DNA was performed using Illumina Miseq, Pacific Biosciences (PacBio), and Oxford Nanopore Technology (ONT) in parallel. The raw sequencing reads were analyzed with Nanoplot [42].

### Genome Assembly from Different Sequencing Methods and Annotation

The genome assembly was performed on the University of Florida high performance computing cluster: Hipergator, using different computing modules. ONT raw reads were trimmed with Porechop (v0.2.4, https://github.com/rrwick/Porechop) and filtered with Filtlong (v0.2.1, -- min_mean_q 90 --min_length 1000, https://github.com/rrwick/Filtlong). Assembly was generated following the workflow of long-reads consensus assembly tool Trycycler (Version: 0.4.2) [44]. Several individual assembly tools were utilized during the initial assembly step, including Flye (Version: 2.9) [113], MinASM (Version: 0.3-r179) [114], Minipolish (Version: 0.1.3) [115], NextDenovo (Version: 2.4.0) [116], NextPolish (Version: 1.3.1) [117] and Raven (Version: 1.8.1) [118]. The consensus assembly was further self-polished with the variant calling and consensus sequence tool medaka (Version: 1.2.1; © 2018-Oxford Nanopore Technologies Ltd.) and polished using Illumina Miseq reads with Pilon (Version: 1.24) [45]. Circularization of the genome was performed using Circlator (Version: 1.5.5) [46]. The assemblies were evaluated by Busco [119] for completeness. The assembled genome was annotated using the NCBI Prokaryotic Genome Annotation Pipeline (PGAP) (https://www.ncbi.nlm.nih.gov/genome/annotation_prok/).

### Phylogenetic Analysis

A phylogenetic analysis was performed using the whole-genome assembly of En-Cren and 321 available whole-genome assemblies with annotated GenBank files of *Enterobacter* strains across all species in the NCBI genome database (as of June 2022). An average nucleotide identity (ANI) analysis was done with the Python program PYANI [47]. Then 32 strains with 95% or more ANI to En-Cren were selected to generate a phylogenetic tree alongside the type strain. Core genome identification of the strains and alignment was conducted using Roary (Version: 3.12.0, setting: - i 90 -e --mafft -cd 100 -g 100000) [120]. RAxML (Version: 8.2.10) [48] was used to create a maximum likelihood tree with 100 bootstrap and GTR gamma model. Illustration of En-Cren genome features and the comparison with closely related strains were performed with web server Circular Genomic Viewer (CGView/Proksee) [121].

### Identification of Prophages and Genomic Islands

Prophage regions in En-Cren were identified using the online tool PHASTER (PHAge Search Tool Enhanced Release) [122], [123]. Genomic island identification was performed with IslandViewer4 [124].

### Targeted Bacterial Mutagenesis

Strains and plasmids used are listed in Table S8. Targeted mutagenesis by plasmid insertion was accomplished by homologous recombination using the pKNOCK-Km vector [125]. The plasmid pKNOCK-Km does not replicate in En-Cren but can replicate high copy numbers in *E.coli* with a *pir* gene [126]. A truncated middle part (300-500bp, around 150bp from the N-terminus) of the targeted gene was amplified by PCR and ligated into the pKNOCK-Km plasmid [125] after restriction enzyme digestion. The plasmid pKNOCK-Km does not replicate in En-Cren but can propagate in high copies in *E. coli* with a *pir* gene mutation [126]. The plasmid construct was transformed into One Shot PIR1 *E. coli* competent cells (Invitrogen™) by heat shock transformation. Plasmid was extracted using Monarch Plasmid Miniprep Kit (New England BioLabs) from the propagated *E. coli* culture and checked for insertion fragments through enzyme digestion and gel electrophoresis. Validated plasmid was then transformed into competent cells of WT En-Cren (prepared from bacterial culture at exponential growth stage with a OD_600_ of 0.5). Transformants were selected by kanamycin (Km) resistance as a sign of successful vector insertion into the bacterial genome and then confirmed for the desired mutation event by PCR product sequencing with a forward primer selected from the En-Cren genome upstream of the targeted gene and a reverse primer selected from the vector sequence downstream of the fragment insertion sites. Genes selected for mutagenesis included the bacterial flagella related genes *(flgE*, *flgD*, *flgF*, *motA*, *flhA*, *filC*, *fliR)*, the type IV pili-related genes (*pilQ*, *pilT*, *pilC, pilA*), and IAA/PAA biosynthesis and catabolite pathway genes (*ipdC, paaK*).

Double recombinant mutants (*flgE-DR*, *flgEFG-DR,* and *island1-* (peg3455-3487-DR)) were also generated with the pKNOCK-Km vector. A gentamycin resistance cassette was flanked by 1 kb sequences upstream and downstream of the target DNA region and inserted into the multiple cloning sites of the plasmid vector. Complementation of a mutant was accomplished by amplifying DNA fragment of the targeted gene with 500bp upstream by PCR and ligating it into the plasmid pHSG396 with chloramphenicol (Cm) resistance cassette. Transformants were selected for pHSG396 vector by screening on media containing chloramphenicol and confirmed by PCR.

Violacein producing mutants were created for colorimetric detection of En-Cren. Plasmid pVIO1-2-Tet was constructed by modifying the plasmid pVIO1-2 [127], which contains the violacein-producing gene cluster *vioABCDE*. A tetracycline resistance cassette was amplified through PCR and ligated into the MluI restriction enzyme site of pVIO1-2 following enzyme digestion (Fig. S5). pVIO1-2-Tet was introduced in wild type and mutant strains of the En-Cren, creating purple-cell bacterial (violacein producing) strains with pVIO1-2-Tet (-pVio).

### Bacterial Growth Rate Measurements

En-Cren WT and mutant strains were incubated in LB medium at 28°C overnight. The overnight cultures were diluted at 1:50 and further incubated to OD_600_= 0.6 as inoculums. 5 ul of these inoculums were transferred to new test tubes with 3 ml LB and shaking incubated at 28°C. OD readings were measured at 600 nm in a GENESYS™ 20 Visible Spectrophotometer (Thermo Scientific) every two hours. In parallel, serial dilution plating and bacterial counts were performed every four hours for 24 hours. Growth curves and an ANOVA test were both done with the plot function, dplyr package [128], ggplot2 package (Wickham, 2009), and ANOVA function [130] in R studio.

### Measurement of IAA and PAA Production

1 ml overnight culture of WT, *ipdC*-mutant, and *ipdC* complemented En-Cren (*ipdC-CP*) in LB broth complemented with antibiotics kanamycin (*ipdC-*) and chloramphenicol (*ipdC-CP*) were inoculated into 5 ml ¼ strength potato dextrose broth (PDB) medium and liquid M9 minimal media [131] using 20% citric acid as carbon source instead of 20% Glucose (MM) supplemented with 0.1% L-tryptophan (Trp) or 0.1% phenylalanine (Phe). These cultures were shaken and incubated for another 24 hours at 28°C, filtered with 0.22 um syringe filters, and the IAA and PAA concentrations were determined. Quantification of the phytohormones IAA and PAA was performed in a method adapted from [132], [133]. Briefly cultures were extracted in 300 µl of H_2_O:1-propanol:HCl (1:2:0.005) spiked with 100 ng of d7-PAA of d5-IAA as internal standards. Samples were homogenized and extracted with 1 ml of methylene chloride. The methylene chloride:1-propanol layer was collected, derivatized with trimethylsilyldiazomethane, quenched with acetic acid in hexane, and volatile compounds were collected by vapor phase extraction. Volatiles were eluted in methylene chloride and analyzed using GC-MS [GC, Agilent 7890B; MS, Agilent 5977B; column, Agilent DB-35MS (30 m; 0.25 mm; 0.25 µm)] with chemical ionization using iso-butane. The derivatized ions monitored are PAA (methyl PAA, m/z 151), d7-PAA (methyl d7-PAA, m/z 158), IAA (methyl IAA, m/z 190) and d5-IAA (methyl d5-IAA, m/z 195). Another group of measurement was performed with En-Cren WT and the *paaK-* mutant with the same method with a 40-hour incubation period after bacteria inoculation instead of 24 hours.

### Co-culture of En-Cren and Fungi

A 6 mm hyphae-agar plug from the freshly growing margin of each fungal culture (Rs-Cren and other collected fungi) was transferred to the center of a new ¼ strength LBA medium plate. After the fungal hyphae on the new ¼ LBA plate grew to the radius of about 8mm (including 3mm of the initial fungal plug), En-Cren strains containing the violacein producing plasmid (pVIO1-2-Tet) were streaked on one side of the same plate about 15mm away from the growing hyphal tips. The co-culture plates were incubated at room temperature for up to two weeks. Pictures of the coculture were taken once every 24 hours afterward to record the fungal growth and bacterial movement patterns. Based on the different fungal/bacterial interactions, the levels of compatibility regarding the hyphal movement of the bacterium on the fungal hyphae were defined as follows: ++ for rapid migration similar to the En-Cren/Rs-Cren, + for positive but slower migration and less bacterial accumulation along the hyphae, - for normal fungal growth but no observable bacterial migration more than one week after contact, -- for inhibition interactions.

### Random mutagenesis of Tn5 transposon mutant Library and Hyphal Movement Mutant Screening

Random mutant libraries of En-Cren bacteria were generated by electroporation with the EZ-Tn5 <KAN-2> Tnp Transposome Kit (Lucigen) and stored in 20% glycerol at -80 °C for future use in the fungal hyphae movement mutant screening. Identification of transposon insertion sites was done by sequencing PCR amplification products using primers for transposon specific sequence and random primers.

Tn5-mutant libraries of En-Cren were used to screen for bacterial mutants that have impaired ability to move along fungal hyphae. A freshly grown hyphae-agar plug was shredded with metal beads and 1 ml of deionized water using a tissue homogenizer. Then each 100 ul of the homogenous fragmented hyphae suspension was spread evenly on a 15 mm ¼ LB agar plate with inoculation loops until dry. The fragmented hyphae plates were incubated at 28°C for 30 hours to let the numerous small fungal colonies establish themselves on the plate. After the growth period, dilutions of the bacterial mutant libraries were adjusted to less than 10^3^ CFU/ml, and 100 ul of the diluted bacterial inoculum was spread on each of the fragmented hyphae plates. These screening plates were incubated at 28°C for another 36 hours before the bacterial colonies that did not move along fungal hyphae were identified and streaked into single colonies as mutant movement candidates for further verification. The candidate mutants were tested using the coculture method described above on individual ¼ LBA plates with at least three replications after being transformed with the violacein-producing plasmid pVIO1-2-Tet.

### Testing of hyphal movement on aerial hyphae

With some of the co-culture plates of En-Cren WT-pVio and Rs-Cren, a thin layer of the ¼ LBA medium was also prepared on the lids of the petri dishes to allow further aerial growth of the fungus and observation of potential En-Cren movement.

### Filming of Time-lapsed Photo Series

A Raspberry Pi Camera was utilized to take photos of cocultured Rs-Cren and violacein producing WT-pvio En-Cren at intervals of 1 hour for 96 hours. The 20-second video was generated by staking the time-lapsed images.

### Bacterial Swimming Motility Assay

Overnight cultures of En-Cren strains in LB liquid medium were diluted to OD_600_= 0.6 as the inoculum. 2 ul of the inoculums were placed at the center of the soft agar (0.3% agar) ¼ LB plates using techniques modified from others [134], [135]. Tryptone agar (0.30% agar) was used in parallel experiments [135]. The swimming phenotypes were examined after 12 hours of incubation at 30°C.

## Acknowledgements

We would like to express sincere gratitude to our colleagues at University of Florida, Dr Jeff Jones, Dr Jeffrey Rollins, and Dr Sixue Chen for their valuable comments and suggestions, and Assistant Scientist Rosanne Healy for providing fungal materials during the research.

Research funding was supported by NSF projects 1258028 and 1741090. Funding for GC-MS analysis in this study was provided by the United States Department of Agriculture-Agriculture Research Service project 6036-11210-002-000-D to AKB.

---

**Fig. S1.**
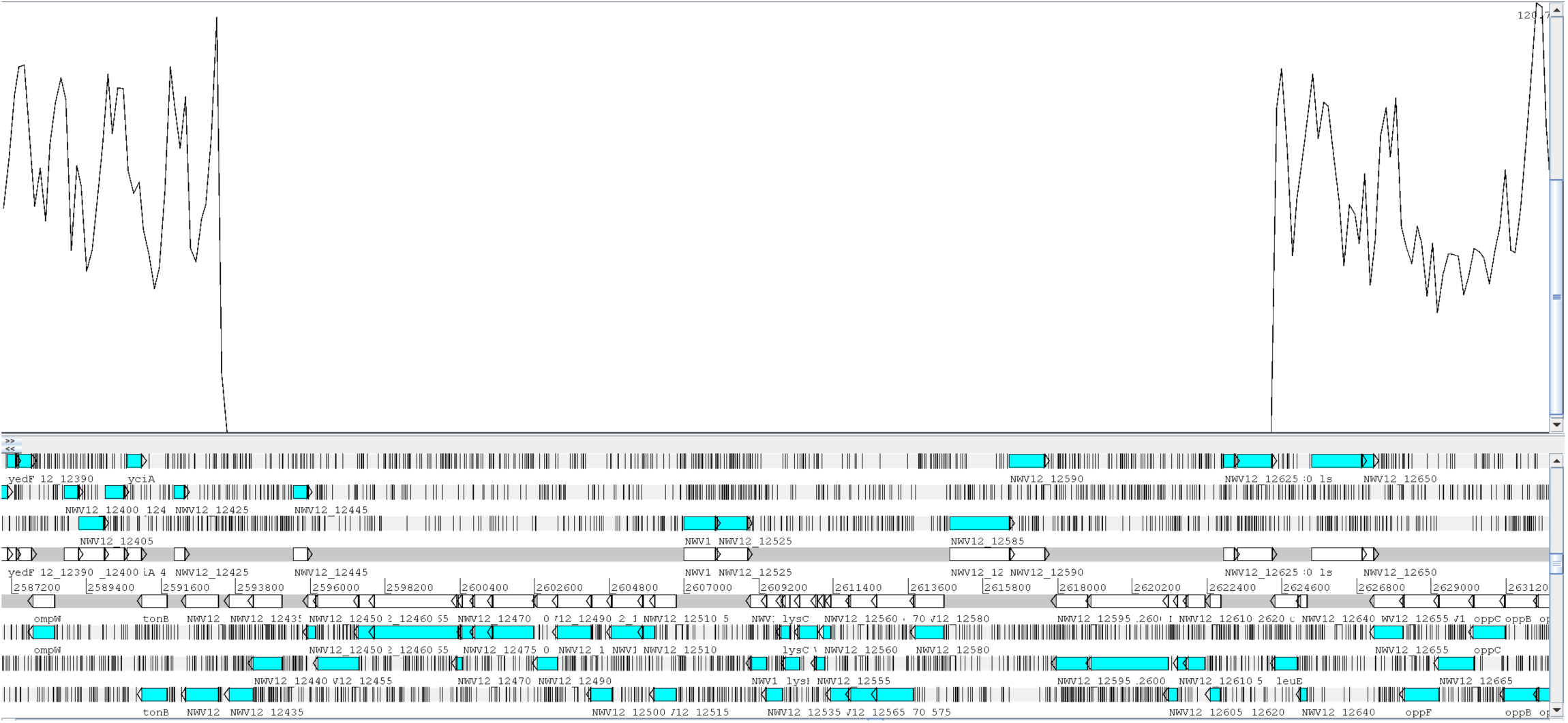
Possible prophage excision revealed when mapping En-Cren *fliR-* mutant illumina sequence reads to WT En-Cren in Artemis. Presumed prophage excision region (prophage region 4) showed no reads coverage, while the general raw reads coverage was at 84x and adjacent reads mapping coverage peaks at around 120x.

**Fig. S2.**
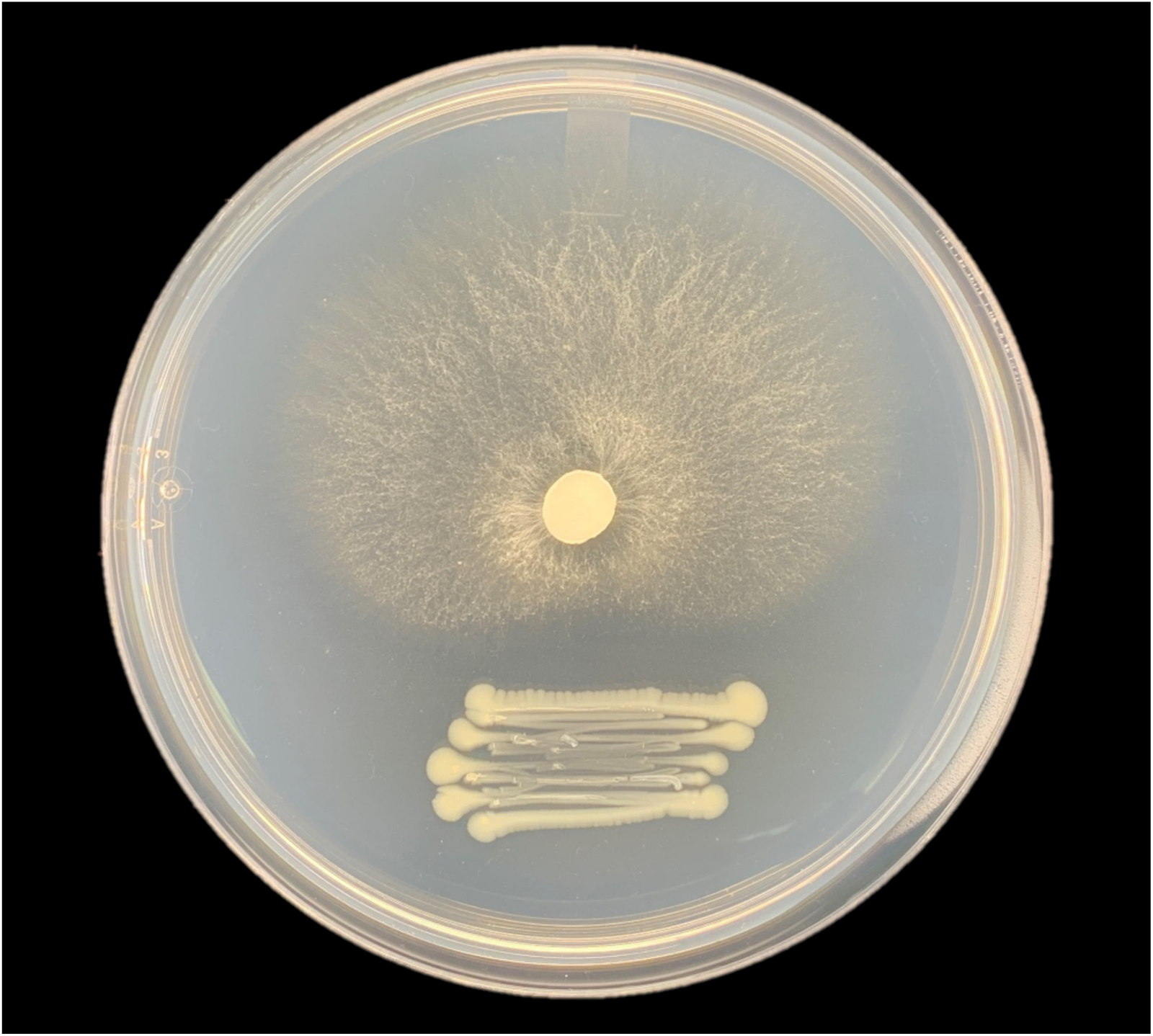
*Ganoderma sessile* co-cultured for 14 days with WT En-Cren (without violacein producing plasmid), showing inhibition of fungal growth by En-Cren without violacein.

**Fig. S3.**
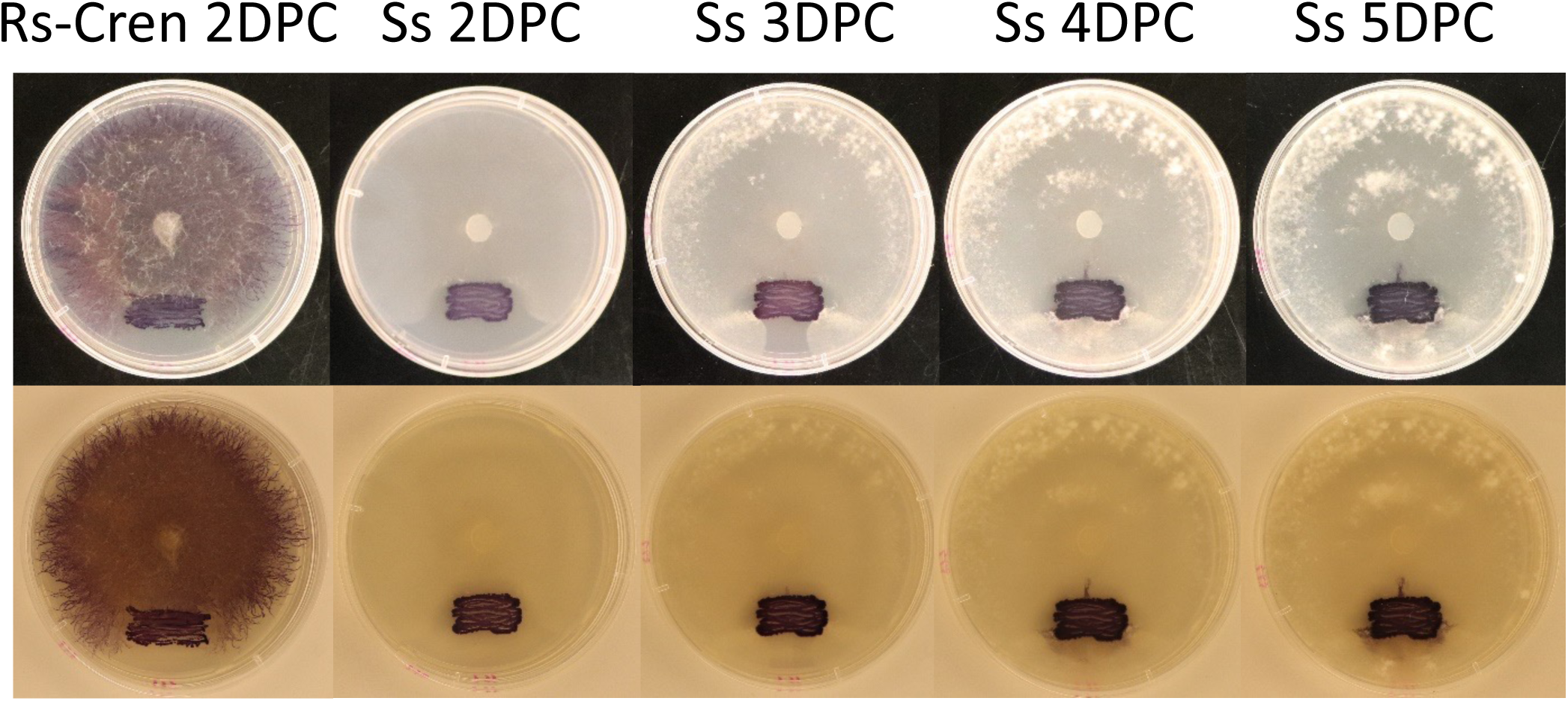
En-Cren movement on a *Sclerotinia sclerotiorum* isolate compared to movement on the original host *R. solani* strain. Picture taken 2, 3, 4 and 5 days after the growing periphery of the fungus reach the bacterial inoculation site.

**Fig. S4.**
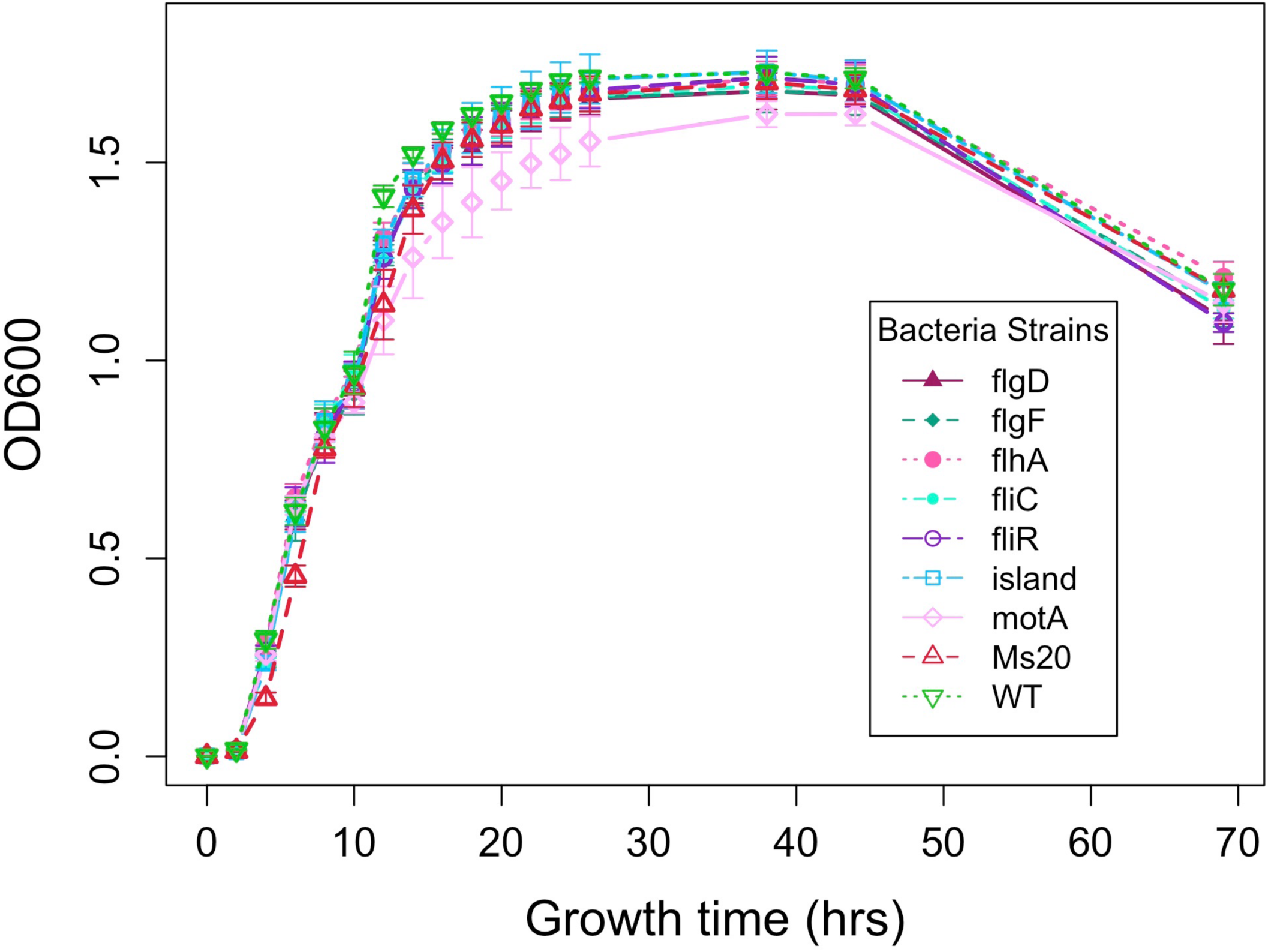
Growth curves of En-Cren WT and motility mutants in LB. Bacterial growth was measured by optical density at 600 nm. ANOVA test has shown no significant difference of the OD600 values between the mutants and the WT strain at each time point.

**Fig. S5.**
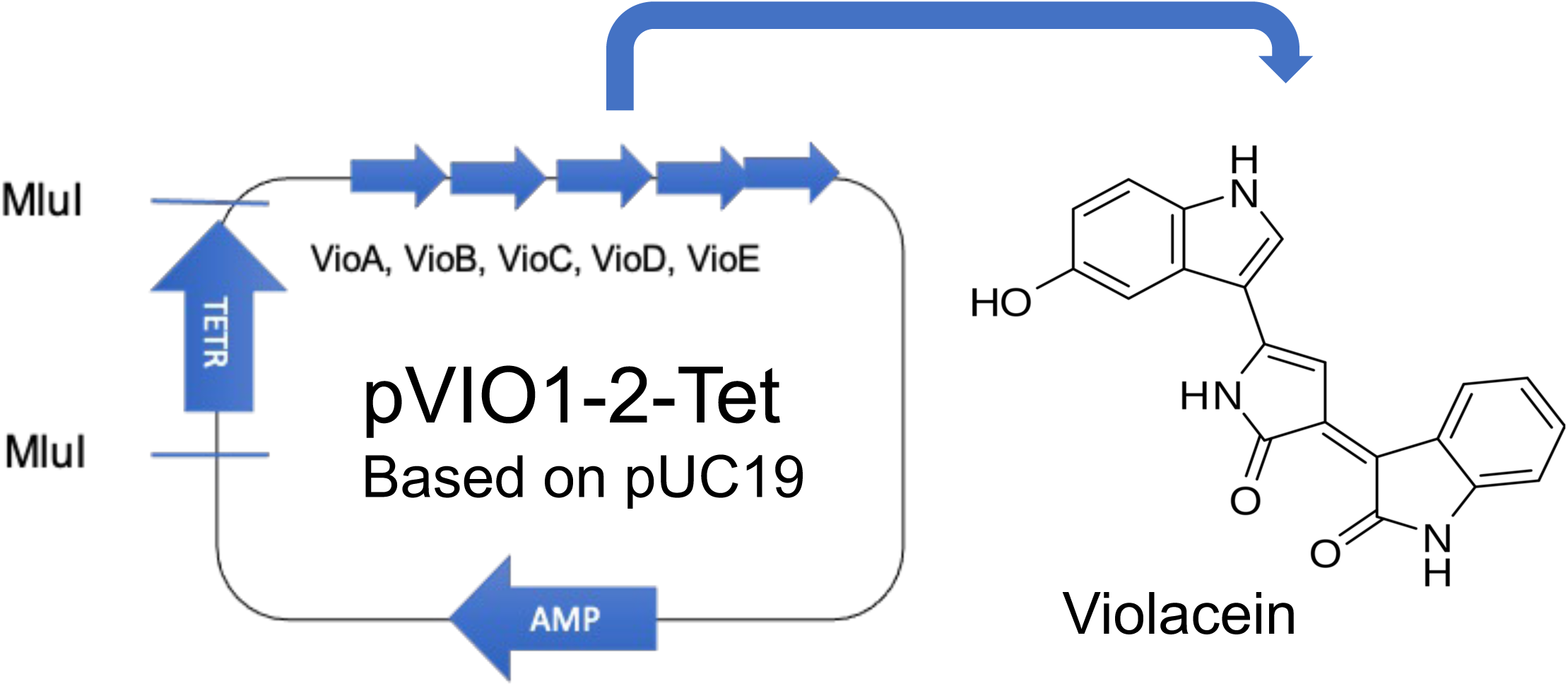
Structure of pVIO1-2 with the violacein producing gene cluster vioABCDE. The plasmid was modified with tetracycline resistance gene insertion at MluI enzyme site.

**Supplement Table S1.**
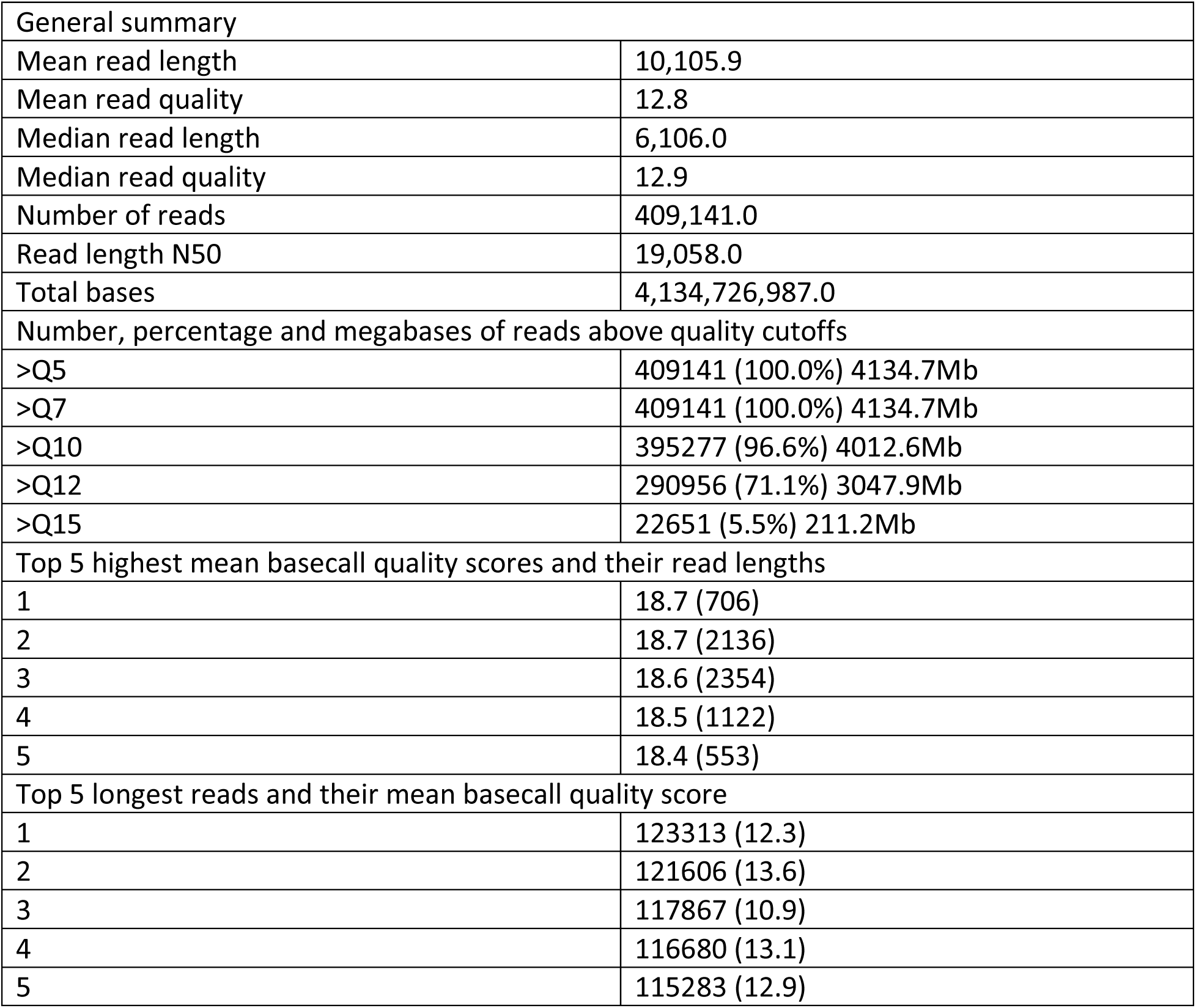
Nanopore sequencing summary.

|  |  |
| --- | --- |
| General summary |  |
| Mean read length | 10,105.9 |
| Mean read quality | 12.8 |
| Median read length | 6,106.0 |
| Median read quality | 12.9 |
| Number of reads | 409,141.0 |
| Read length N50 | 19,058.0 |
| Total bases | 4,134,726,987.0 |
| Number, percentage and megabases of reads above quality cutoffs |  |
| >Q5 | 409141 (100.0%) 4134.7Mb |
| >Q7 | 409141 (100.0%) 4134.7Mb |
| >Q10 | 395277 (96.6%) 4012.6Mb |
| >Q12 | 290956 (71.1%) 3047.9Mb |
| >Q15 | 22651 (5.5%) 211.2Mb |
| Top 5 highest mean basecall quality scores and their read lengths |  |
| 1 | 18.7 (706) |
| 2 | 18.7 (2136) |
| 3 | 18.6 (2354) |
| 4 | 18.5 (1122) |
| 5 | 18.4 (553) |
| Top 5 longest reads and their mean basecall quality score |  |
| 1 | 123313 (12.3) |
| 2 | 121606 (13.6) |
| 3 | 117867 (10.9) |
| 4 | 116680 (13.1) |
| 5 | 115283 (12.9) |

**Supplement Table S2:** Illumina sequencing summary.

| Raw Data | Illumina_R1 | Illumina_R2 |
| --- | --- | --- |
| Mean read length (bp) | 234.4 | 231.2 |
| Mean read quality | 34.3 | 29.1 |
| Median read length (bp) | 249.0 | 248.0 |
| Median read quality | 37.3 | 29.2 |
| Number of reads | 970,457 | 970,457 |
| Read length N50 (bp) | 249.0 | 249.0 |
| Total bases (bp) | 227,451,386 | 224,340,164 |
| Sequences flagged as poor quality | 0 | 0 |
| %GC | 56 | 56 |
| Raw coverage | 49x | 48x |
| Number of reads after Q30 filter | 954,051 | 954,051 |

**Supplement Table S3:** Pangenome statistics of En-Cren with thirty-two closely related strains and type strain type-strain of *Enterobacter*, *Enterobacter cloacae* subsp. *cloacae* ATCC 13047 using Roary.

|  |  |
| --- | --- |
| Core genes (100% <= strains <= 100%) | 2982 |
| Soft core genes (95% <= strains < 100%) | 305 |
| Shell genes (15% <= strains < 95%) | 1693 |
| Cloud genes (0% <= strains < 15%) | 11859 |
| Total genes (0% <= strains <= 100%) | 16839 |

**Supplement Table S4:**
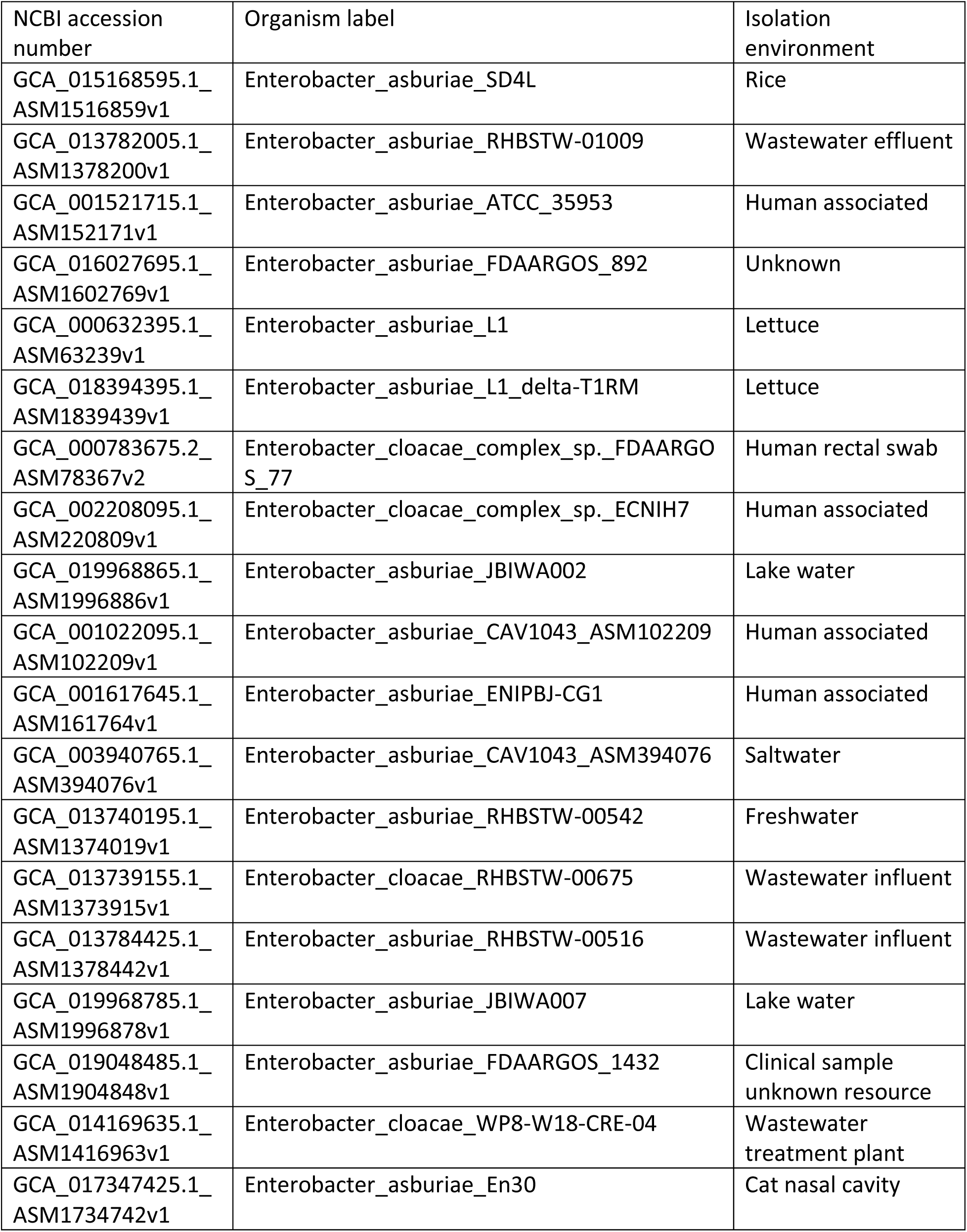
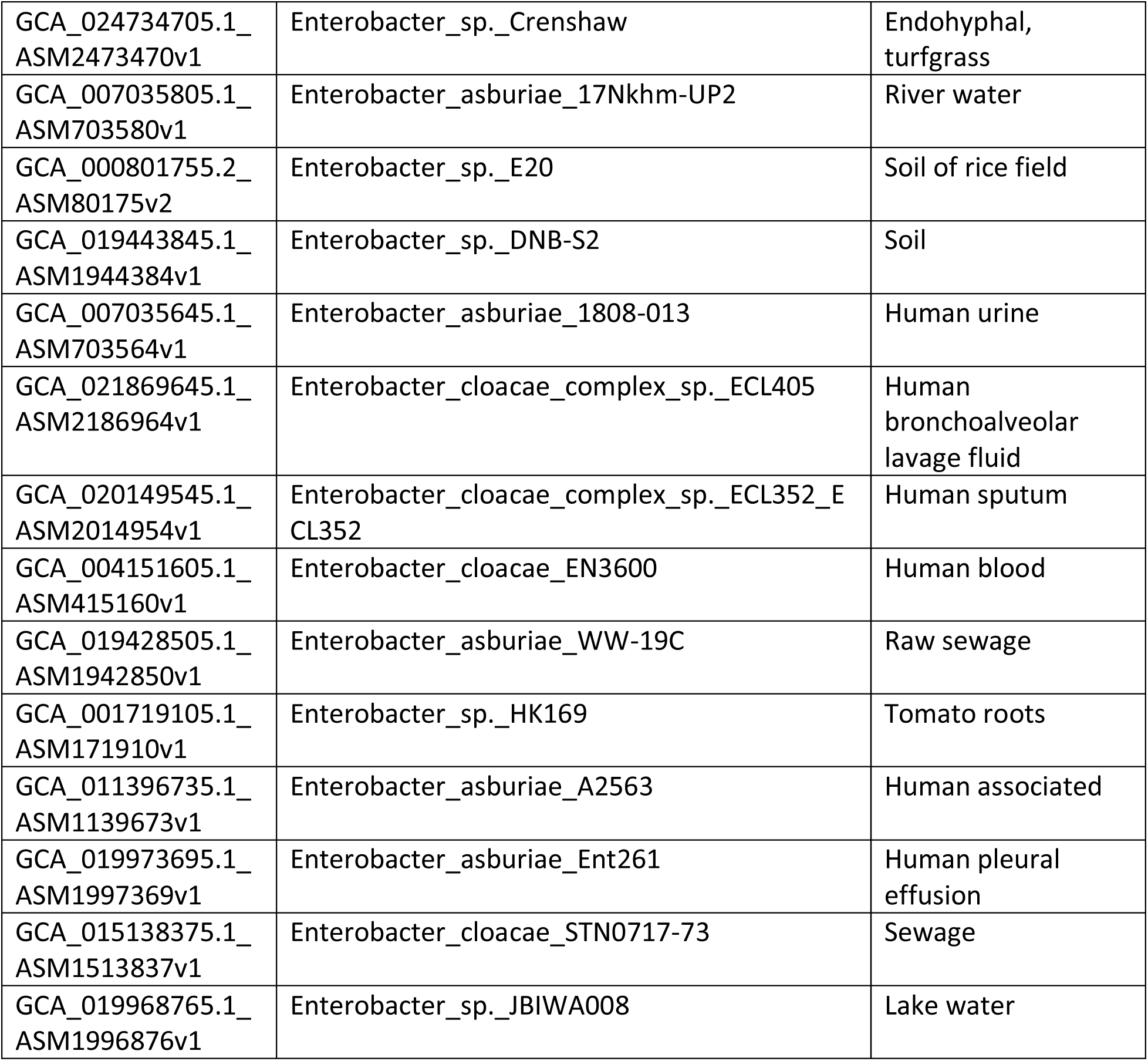
List of En-Cren closely related strains included in the phylogenetic analysis, with corresponding NCBI accession number and origin of isolation.

**Supplement Table S5:**
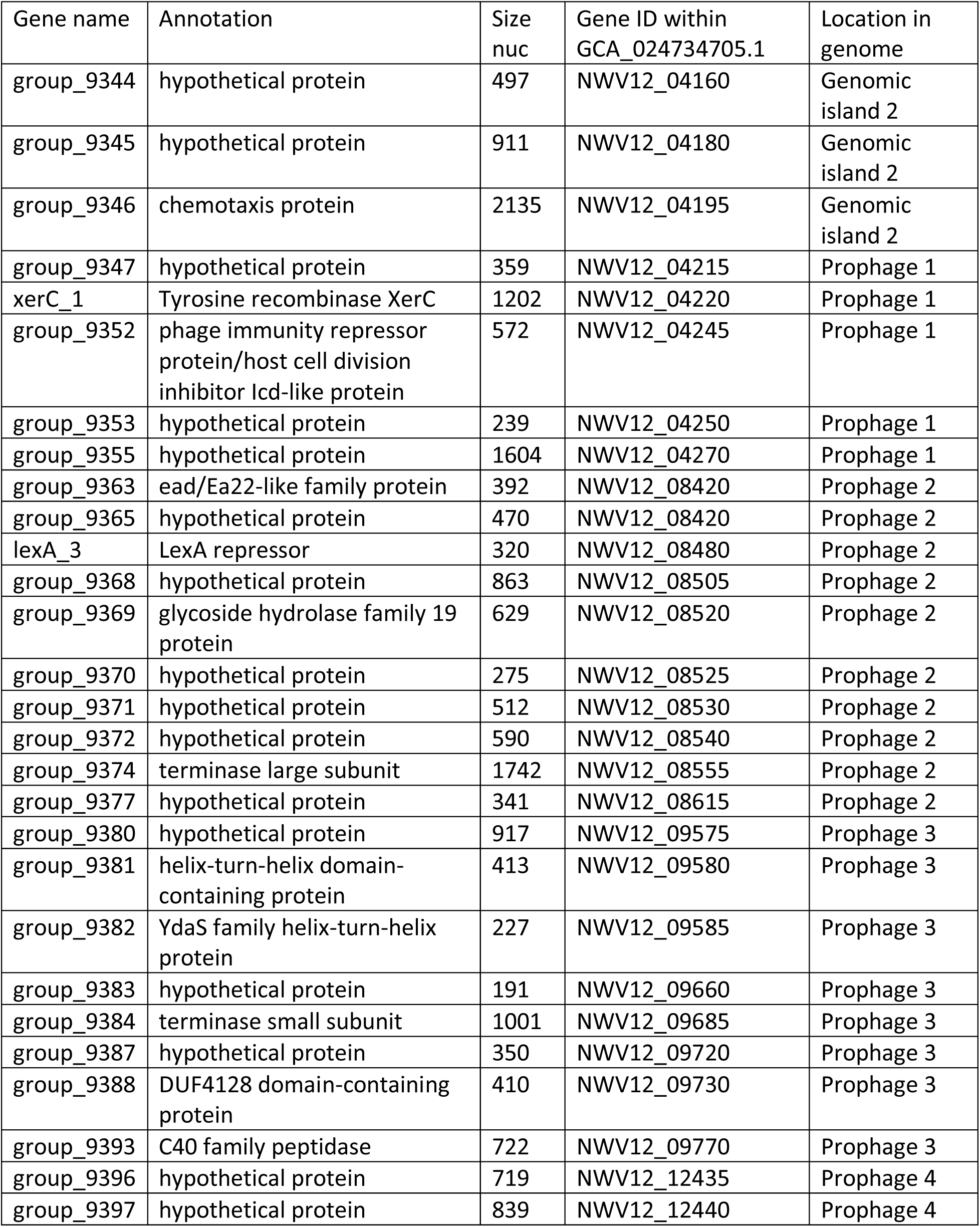
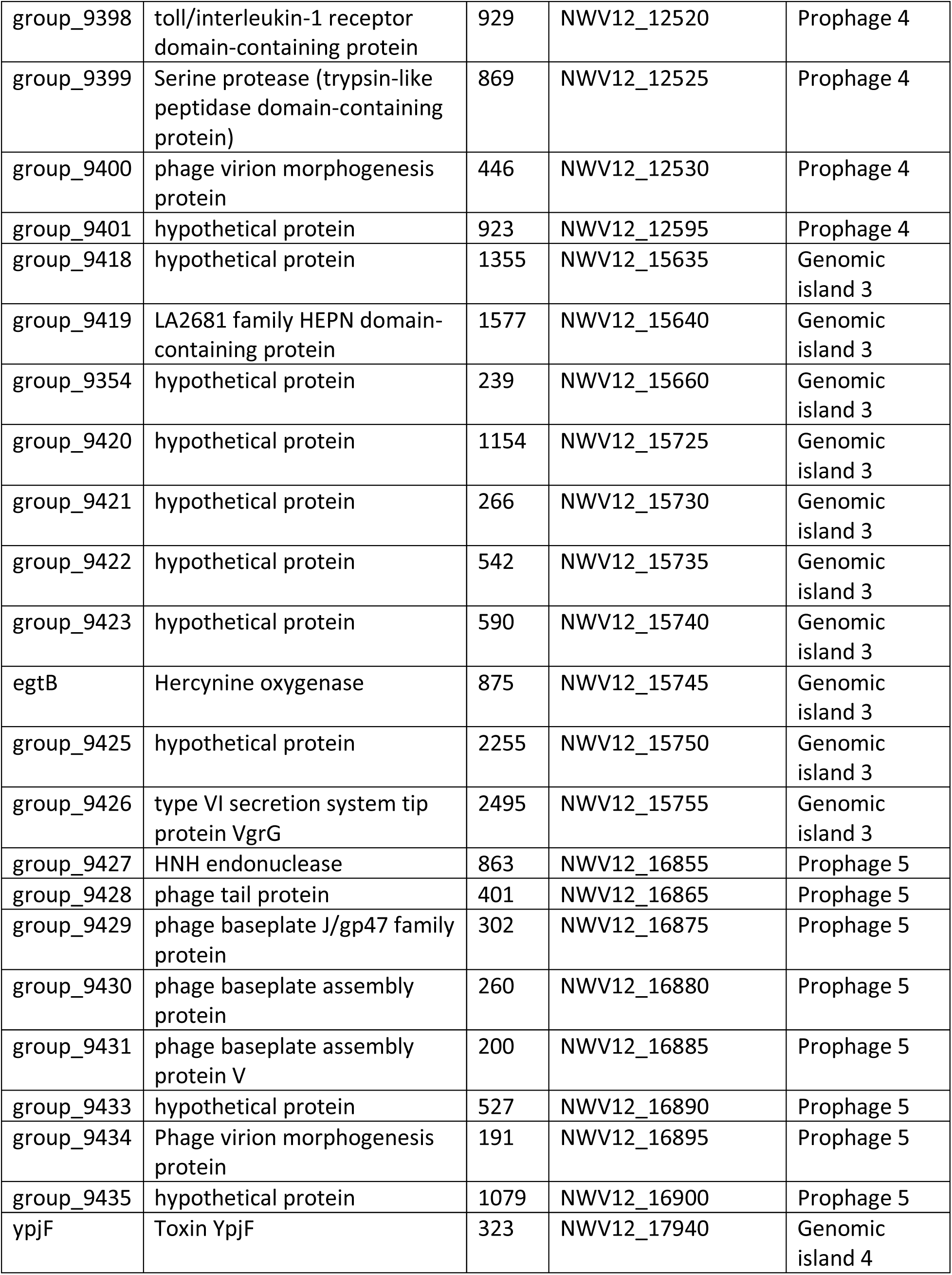
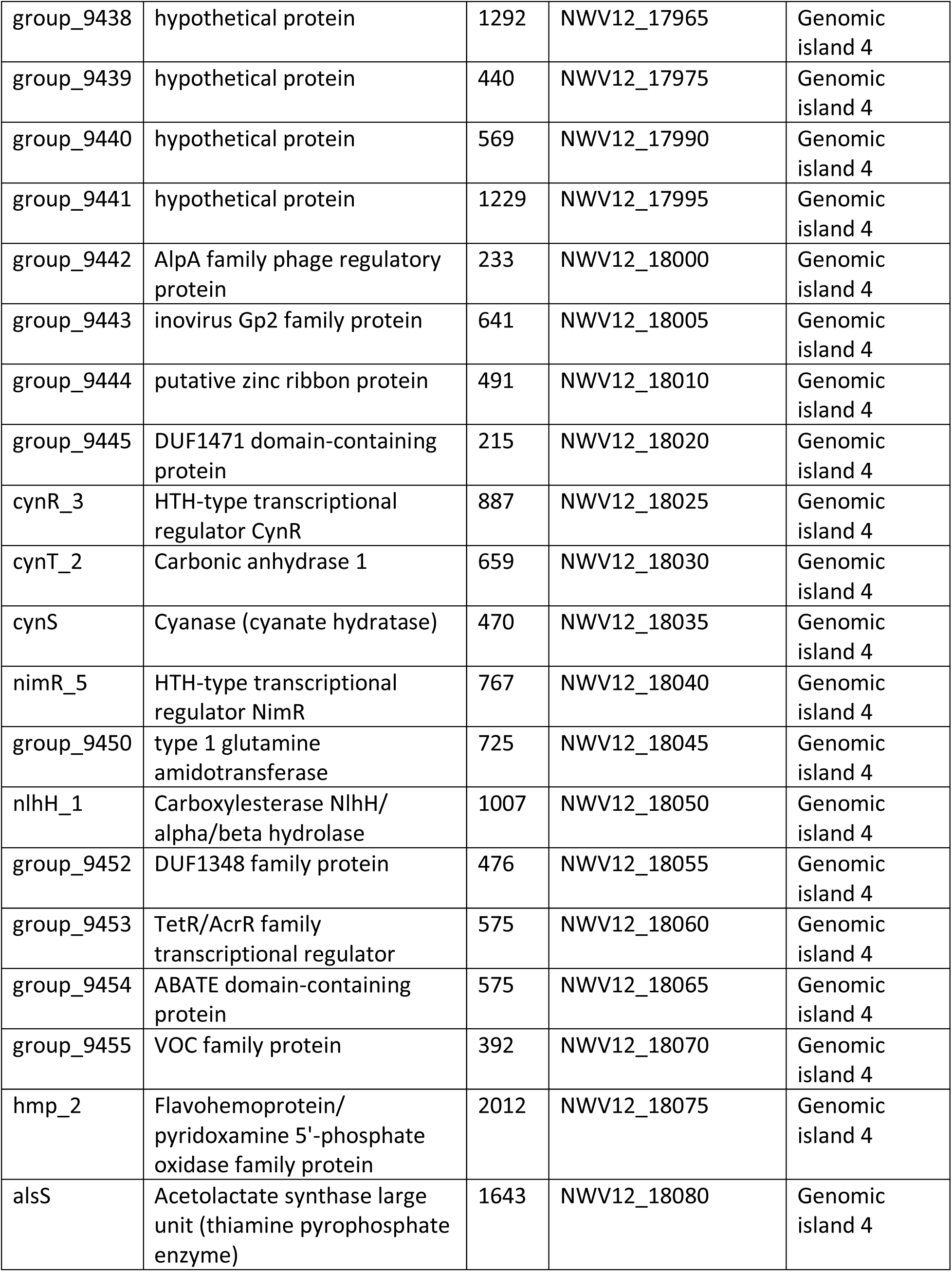
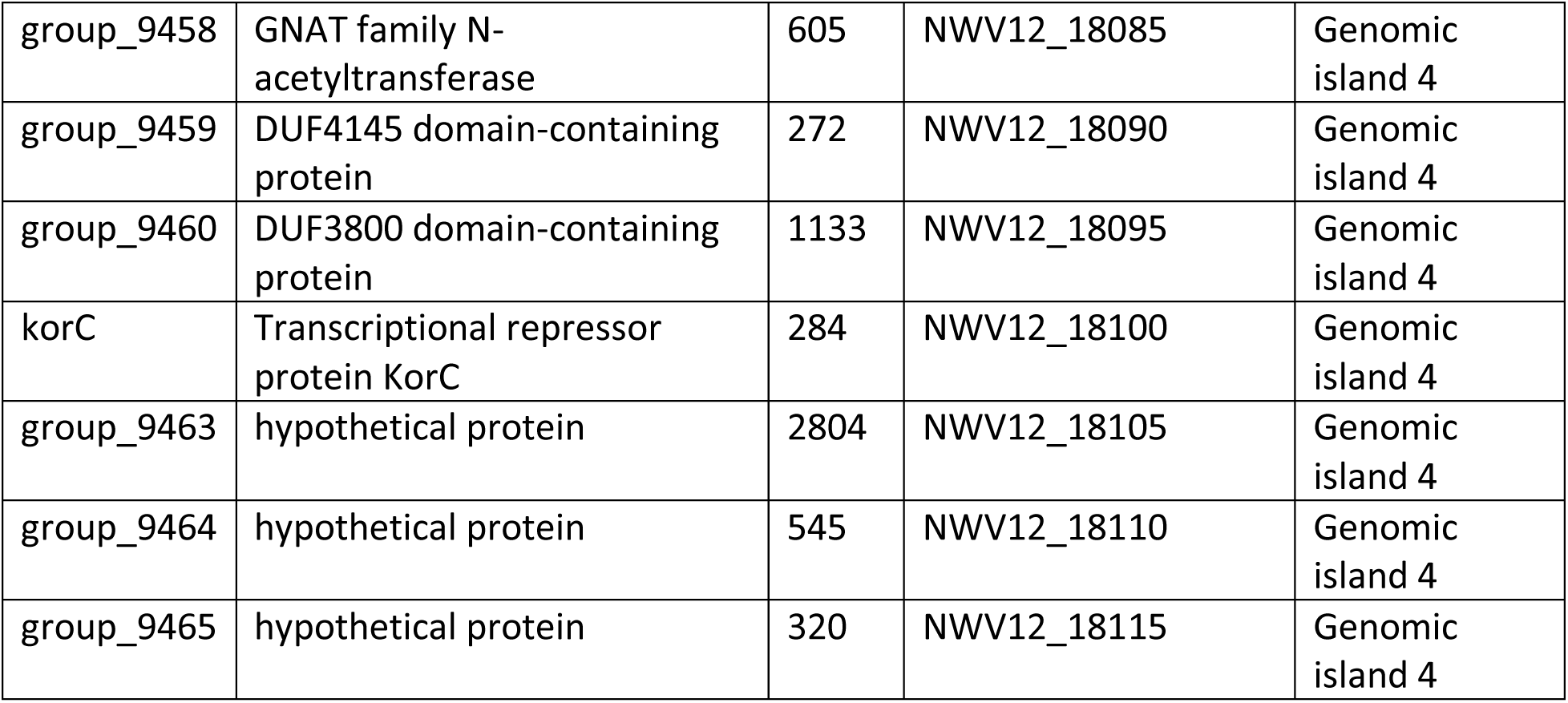
List of En-Cren unique gene with nucleotide size and annotation from Prokka and through NCBI blastP against the non-redundant protein database.

**Supplement Table S6:**
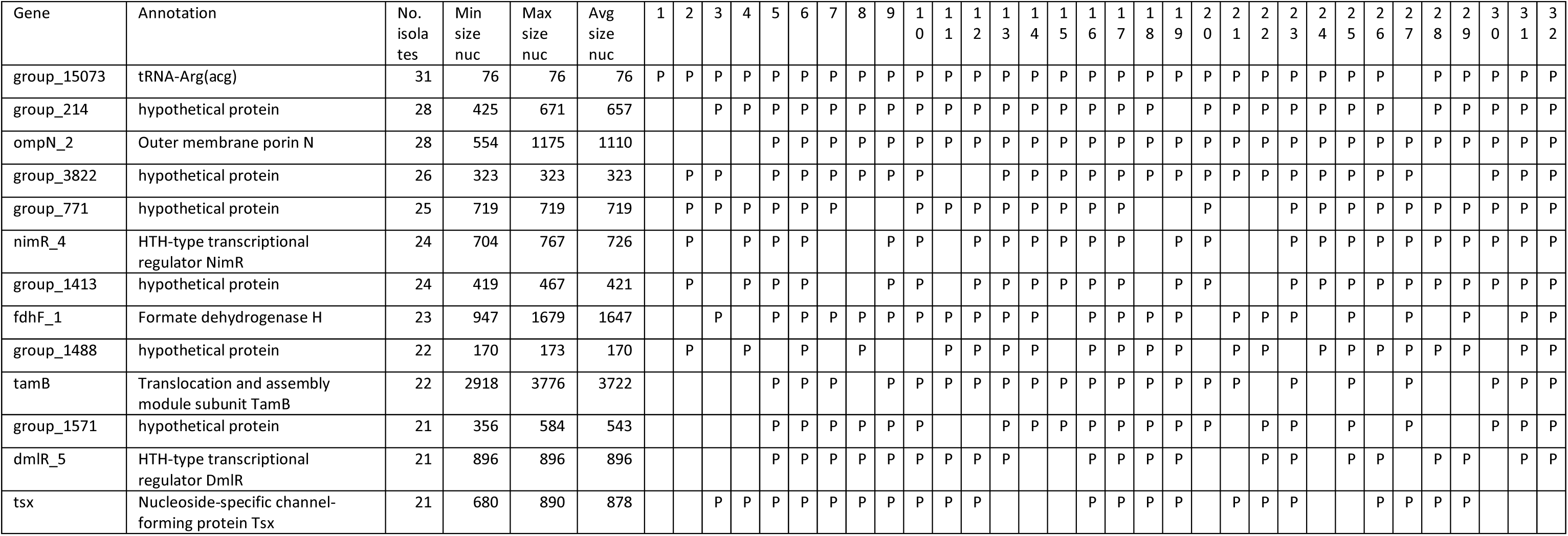
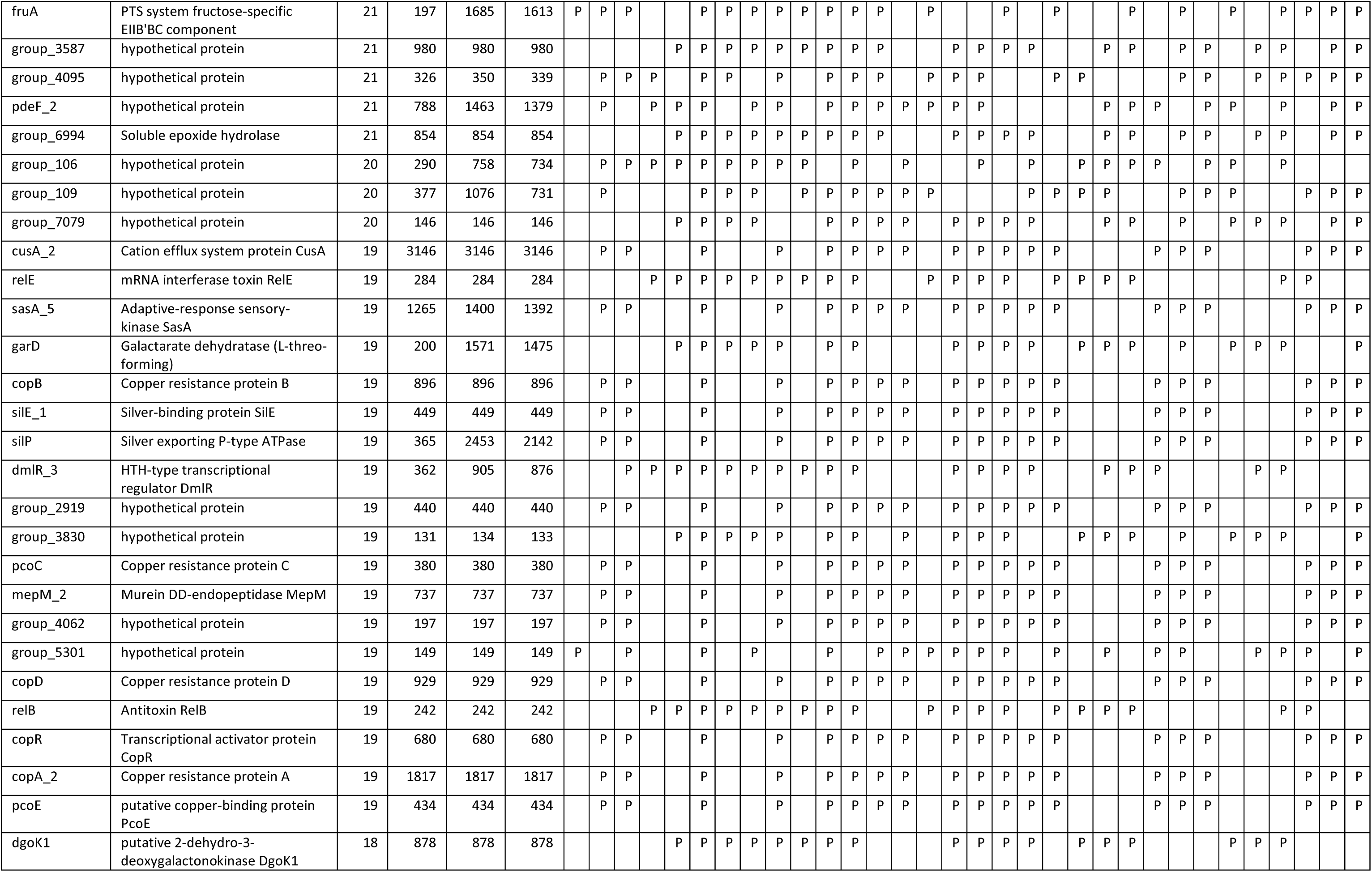
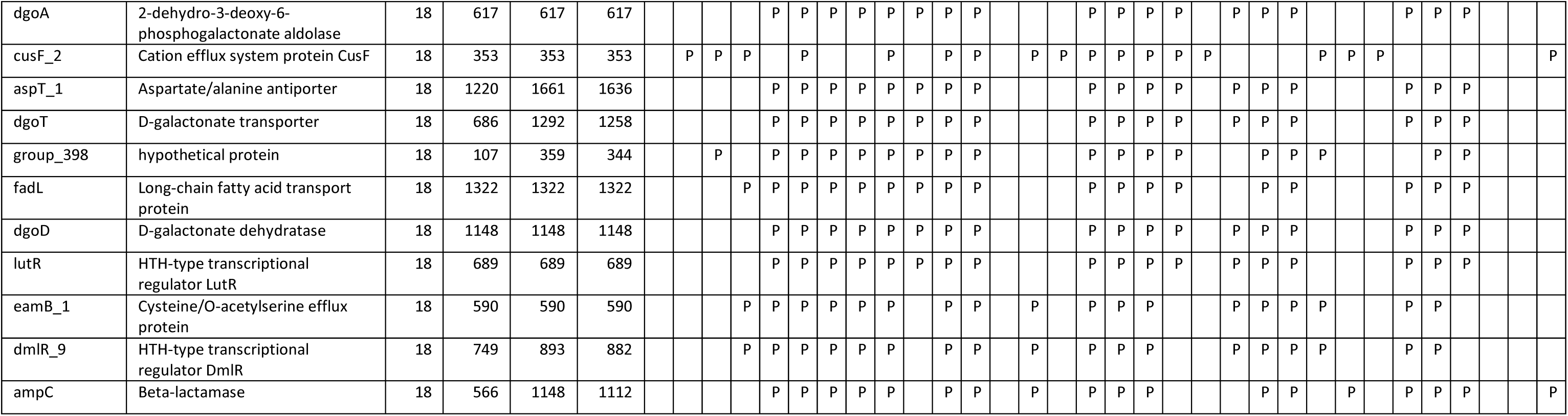
Genes absent in En-Cren genome but present in at least half of the other thirty-two closely related strains according to Roary analysis are listed with annotation and their nucleotide size. Strains are number labeled as follows: 1: GCA_007035805.1_ASM703580v1, 2: GCA_017347425.1_ASM1734742v1, 3: GCA_014169635.1_ASM1416963v1, 4: GCA_000801755.2_ASM80175v2, 5: GCA_000632395.1_ASM63239v1, 6: GCA_000783675.2_ASM78367v2, 7: GCA_001022095.1_ASM102209v1, 8: GCA_001521715.1_ASM152171v1, 9: GCA_001617645.1_ASM161764v1, 10: GCA_001719105.1_ASM171910v1, 11: GCA_002208095.1_ASM220809v1, 12: GCA_003940765.1_ASM394076v1, 13: GCA_004151605.1_ASM415160v1, 14: GCA_007035645.1_ASM703564v1, 15: GCA_011396735.1_ASM1139673v1, 16: GCA_013739155.1_ASM1373915v1, 17: GCA_013740195.1_ASM1374019v1, 18: GCA_013782005.1_ASM1378200v1, 19: GCA_013784425.1_ASM1378442v1, 20: GCA_015138375.1_ASM1513837v1, 21: GCA_015168595.1_ASM1516859v1, 22: GCA_016027695.1_ASM1602769v1, 23: GCA_018394395.1_ASM1839439v1, 24: GCA_019048485.1_ASM1904848v1, 25: GCA_019428505.1_ASM1942850v1, 26: GCA_019443845.1_ASM1944384v1, 27: GCA_019968765.1_ASM1996876v1, 28: GCA_019968785.1_ASM1996878v1, 29: GCA_019968865.1_ASM1996886v1, 30: GCA_019973695.1_ASM1997369v1, 31: GCA_020149545.1_ASM2014954v1, 32: GCA_021869645.1_ASM2186964v1 . “P” refers to presence in the genome assembly.

**Supplement Table S7.**
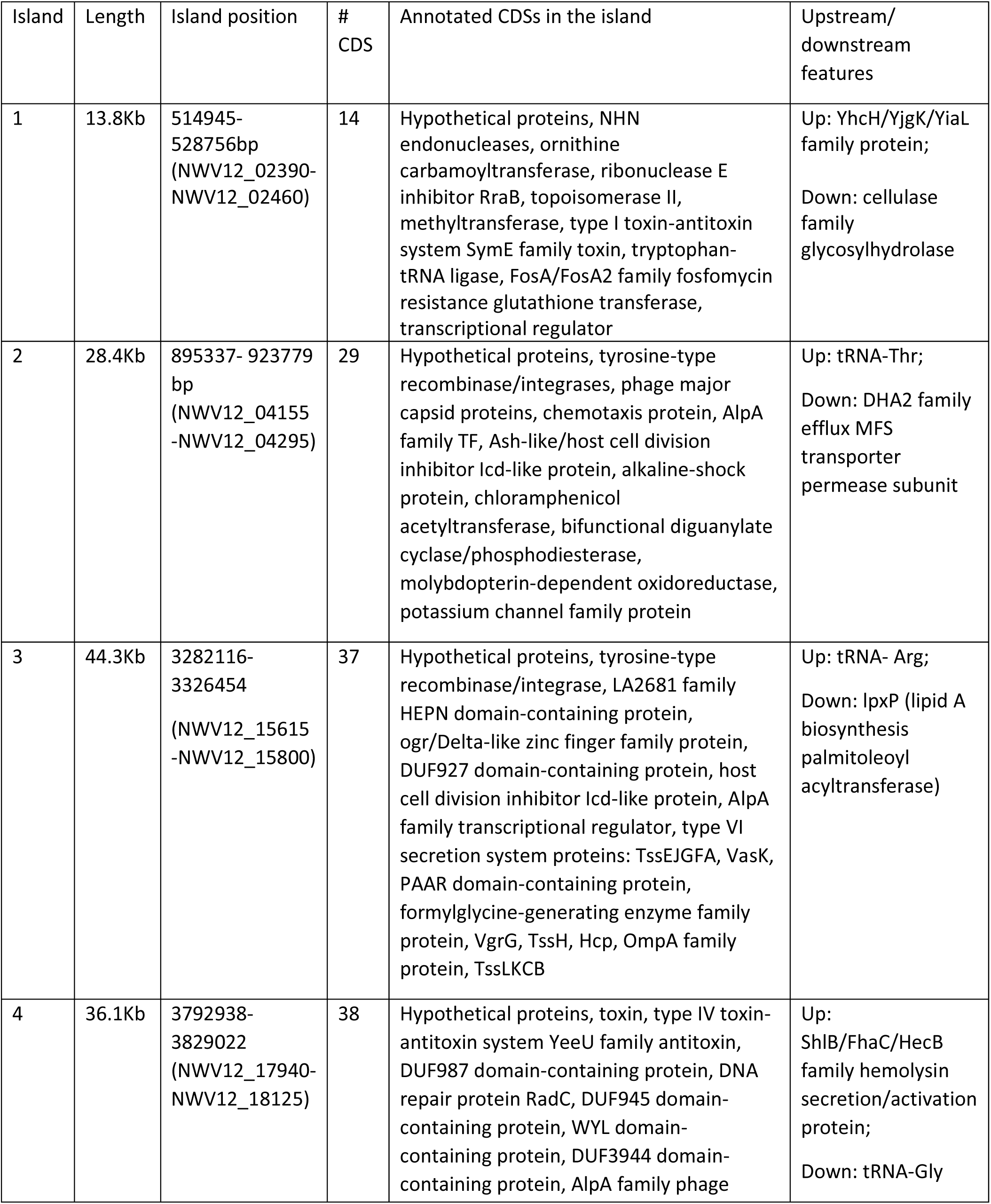
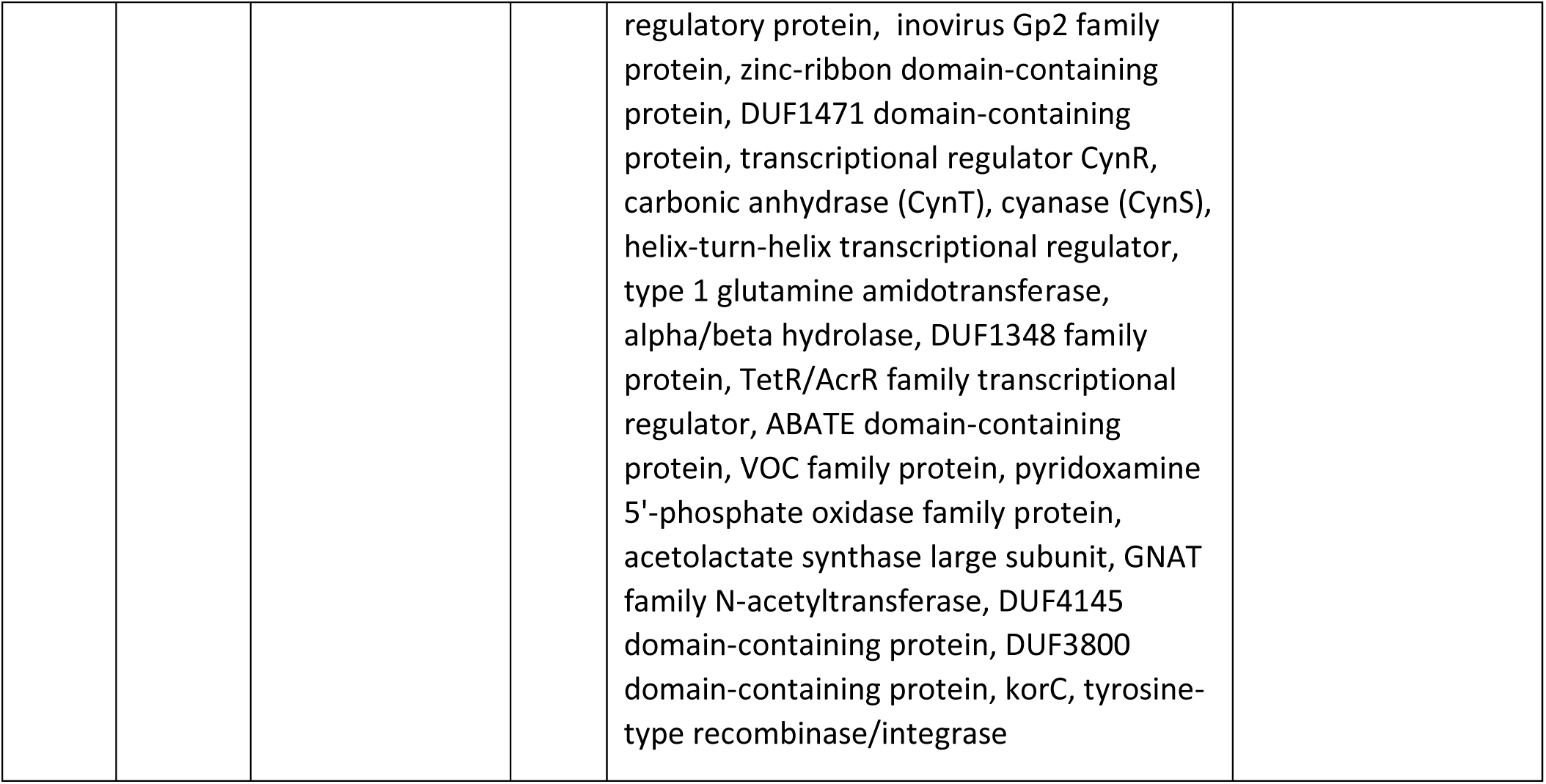
The size, position and list of genes in the unique genomic islands.

**Supplement Table S8.** Summary of En-Cren mutants, their swimming motility and hyphal motility. ‘++’ refers to similar speed and pattern as WT EnCren; ‘+’ refers to slower movement that eventually reach the similar spread as WT EnCren; ‘+/-’ refers to local and limited movement; ‘-’ refers to depletion of visible motility during the time frame of the experiment.

| Mutants | Corresponding Gene product | Method | Swimming motility | Hyphal movement |
| --- | --- | --- | --- | --- |
| *flgE*- | Flagellar hook protein | pKNOCK-Km insertion | - | - |
| *flgF*- | Flagellar basal-body rod protein | pKNOCK-Km insertion | - | - |
| *flgD*- | Basal-body rod modification protein | pKNOCK-Km insertion | - | - |
| *fliR*- | Flagellar biosynthetic protein | pKNOCK-Km insertion/<br>Tn5 random insertion | - | - |
| *fliC*- | Flagellin | pKNOCK-Km insertion | - | - |
| *flhA*- | Flagellar biosynthesis protein | pKNOCK-Km insertion | - | - |
| *flhC*- | Flagellar transcriptional regulator | Tn5 random insertion | - | - |
| *flhD*- | Flagellar transcriptional regulator | Tn5 random insertion | - | - |
| *motA*- | stator element of the flagellar motor | pKNOCK-Km insertion | - | - |
| *pilQ*- | Type IV pilus secretin | pKNOCK-Km insertion | ++ | ++ |
| *pilT*- | Twitching motility protein (Type IV pilus retraction ATPase) | pKNOCK-Km insertion | + | + |
| *pilC*- | Type IV pilus assembly protein | pKNOCK-Km insertion | ++ | ++ |
| *pilA*- | Type IV major pilin protein | pKNOCK-Km insertion | ++ | ++ |
| *pdhR*- | Pyruvate dehydrogenase complex repressor | Tn5 random insertion | + | +/- |

**Supplement Table S9.** List of the collection of fungi with light-colored mycelium like Rs-Cren.

| Fungal strain | Phylum | Culture medium |
| --- | --- | --- |
| *Ganoderma sessile* lqqmo | Basidiomycetes | PDA |
| *Ceratorhiza* sp. 5-15B | Basidiomycetes | PDA |
| *Rhizoctonia solani* | Basidiomycetes | APDA |
| *Cyathus* sp. | Basidiomycetes | PDA |
| *Geotrichum candidum* | Ascomycetes | APDA |
| *Fusarium fujikurai* | Ascomycetes | PDA |
| *Pestalotiopsis* sp. | Ascomycetes | APDA |

